# Diverse *Pseudomonas* species engage in beneficial and suppressive interactions with the kiwifruit pathogen *Psa* across *Actinidia* germplasm

**DOI:** 10.1101/2025.01.31.635981

**Authors:** Haileigh R. Patterson, Lauren M. Hemara, Matthew D. Templeton, Jay Jayaraman

## Abstract

In 2010, a *Pseudomonas syringae* pv. *actinidiae* biovar 3 (Psa3) incursion into New Zealand kiwifruit orchards devastated susceptible *Actinidia chinensis* cultivars. In contrast, many *Actinidia* species maintained in germplasm collections were resistant to Psa3 and showed limited symptoms. Recent genome biosurveillance revealed the emergence of widespread leaf spot symptoms in Psa3-resistant *Actinidia* germplasm. Surprisingly, few Psa3 isolates were recovered from symptomatic tissues, despite the frequent isolation of phenotypical *Pseudomonas* isolates on selective agar. Despite the poor recovery of Psa3 isolates, Psa3 was found in all symptomatic leaf tissue through qPCR and metabarcoding analysis. Metabarcoding revealed stark differences in bacterial community composition from samples taken from lesion-carrying or lesion-free material from the same leaf. Host genotype also appeared to influence community composition but to a lesser extent. Whole genome sequencing of diverse *Pseudomonas* spp. isolates revealed that many belonged to the *P. syringae* species complex. Curiously, the kiwifruit-associated *P. syringae* pv. *actinidifoliorum* (Pfm), was never recovered, nor were any other phylogroup 1 pathovars. Pathogenicity and competitive assays revealed that while individually diverse *Pseudomonas* isolates were not more pathogenic than Pfm or Psa3 on resistant *Actinidia* hosts, they could interact with Psa to sometimes improve their growth, suggesting that these isolates may form pathogenic consortia in these disease-associated phyllosphere communities on typically Psa3-resistant hosts.

**Graphical Abstract:** 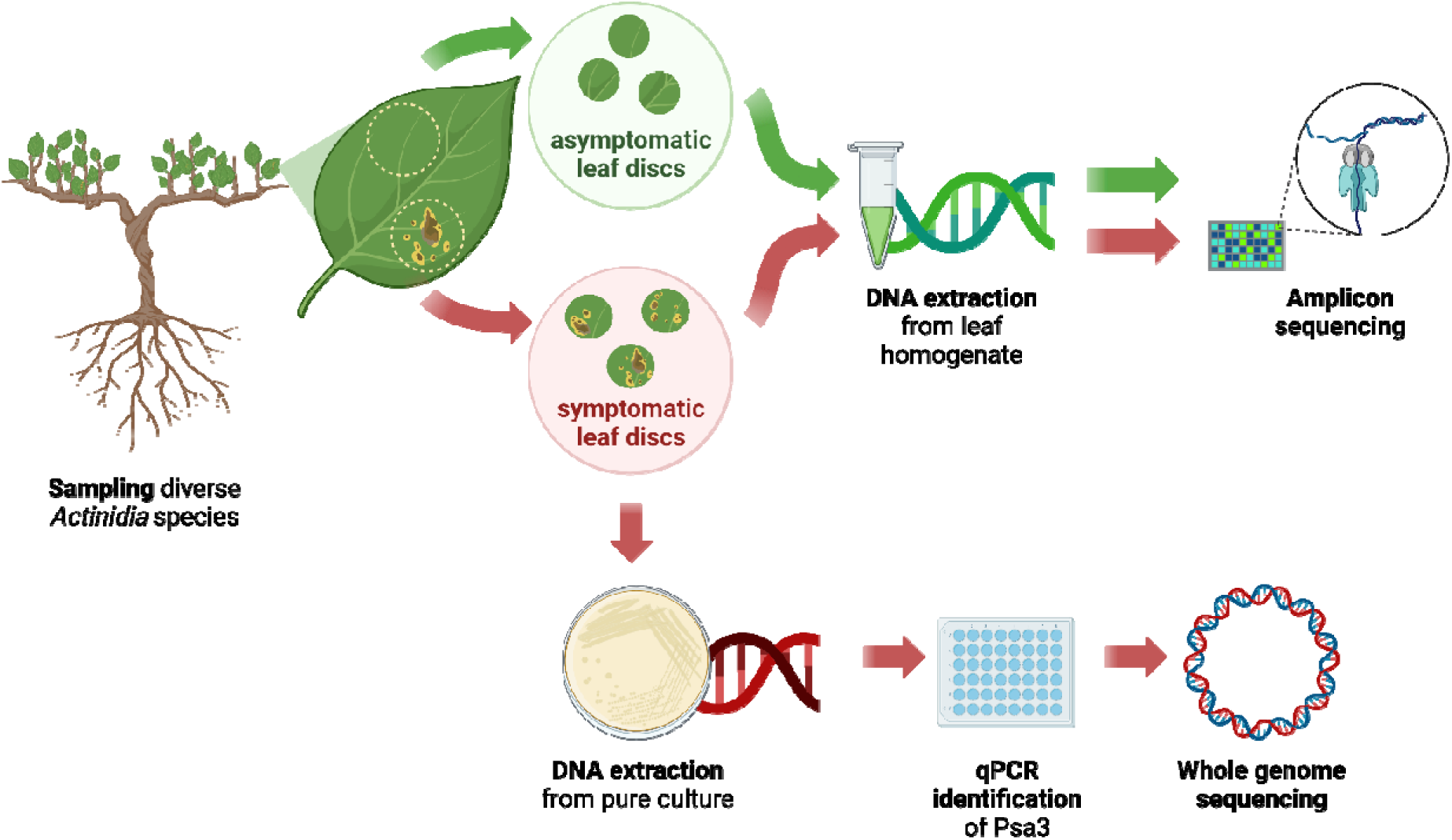

## Introduction

The phyllosphere is one of the most abundant and diverse habitats in the world, host to commensal and mutualistic epiphytes (Agbavor et al., 2022), as well as phytopathogens transitioning from epiphytic to endophytic lifestyles (Vorholt, 2012). Bacteria are the most abundant inhabitants of the phyllosphere (Leveau, 2015; Peñuelas & Terradas, 2014), although the leaf surface is also occupied by a wide range of prokaryotic and eukaryotic microorganisms (Sohrabi et al., 2023). The phyllosphere differs from the widely studied rhizosphere, as each leaf is exposed to the environment, where colonisation can be short-lived and adaptations to harsh conditions are paramount (Lindow & Brandl, 2003; Vorholt, 2012; Koskella, 2023). Further still, each leaf is a microclimate that experiences temperature, water, light, and humidity fluctuations, punctuated by plant architecture of trichomes and stomata (Vacher et al., 2016). Phyllosphere bacteria, therefore, often exist in multi-species aggregates to protect themselves from harsh conditions on the leaf surface (Vorholt, 2012). The exposure of the leaf to the environment not only selects for certain microbes, but also means the possibilities for bacterial introduction are greater, through both endophytic and external colonisation pathways such as the seed and rhizosphere microbiome, biological vectors including insects, and abiotic factors like wind and rain (Hacquard et al., 2017).

Bacteria in the phyllosphere can be considered through two lenses: the way they interact with the plants they live on (host-microbe interactions), and how they interact with other inhabitants of the phyllosphere (microbe-microbe interactions). Within host-microbe interactions, bacteria exist on a spectrum of mutualism to pathogenicity, along which they may transition (Cook et al., 2015; Drew et al., 2021). Regardless of where bacteria sit on the symbiosis spectrum, they share common strategies for symbiosis, including motility, chemotaxis, metabolic adaptation for both competition and host immune suppression, and environmental response (Wiesmann et al., 2023). Much like host-microbe interactions, microbe-microbe interactions can also exist on a spectrum of interactions, as co-virulence, opportunism, and suppression may occur simultaneously within the phyllosphere. For example, phyllosphere bacteria can form pathogenic consortia, altering disease severity and bacterial growth dynamics (Sadhukhan et al., 2024). In the field, the intersection of host-microbe and microbe-microbe interactions is likely to be complex and, to date, is poorly understood. However, recent work has begun to elucidate the key roles and mechanisms of the phyllosphere microbiome in disease (Purahong et al., 2018; Su et al., 2024).

*Pseudomonas syringae* is one of the most widely studied phyllosphere bacteria owing to its ubiquitous and often pathogenic nature (Monteil et al., 2016). In particular, members of the *P. syringae* species complex cause disease more frequently than any other bacterial species, often implicated in new epidemics and posing a major threat to food production globally (Morris et al., 2019). Like other phytopathogens, *P. syringae’s* virulence strategy includes the secretion of effectors and toxins into host cells, which facilitate pathogen entry into the host, suppress host defences, promote pathogen proliferation in host tissue, and aid in nutrient acquisition (Lee et al., 2012). However, many *P. syringae* also have beneficial or neutral interactions with plant hosts and can be found ubiquitously in the environment as part of the water cycle (Morris et al., 2008). Additionally, closely related strains may also be on opposite ends of this spectrum, such as commensal *P. fluorescens* isolates that protect *Arabidopsis thaliana* from genetically similar pathogenic isolates (Wang et al., 2022). This makes *P. syringae* particularly interesting for study, as they represent both established and potential pathogens, allowing us to decipher what may allow new incursions, and the evolution of pathogenesis.

In 2010, *P. syringae* pv. *actinidiae* biovar 3 (Psa3) was isolated from kiwifruit vines in New Zealand (Everett et al., 2011). In susceptible hosts, Psa3 causes necrotic leaf spots, cane canker, vine dieback, bud rot and flower wilting, sometimes leading to the loss of entire vines. Psa3 caused significant financial losses and changed the landscape of kiwifruit production. In the years since this global Psa pandemic, extensive research has been undertaken to understand the origins and evolution of the economically important pathogen (McCann et al., 2013, 2017; Colombi et al., 2017; McAtee et al., 2018; Zhao et al., 2019; Donati et al., 2020; Jayaraman et al., 2020; Hemara et al., 2022; Jayaraman et al., 2023; Ishiga et al., 2023; Vlková-Žlebková et al., 2024; Hemara et al., 2024). Despite considerable contemporary understanding of Psa, knowledge about the larger context and communities in which it exists is only now being revealed. For instance, several *Pseudomonas* spp. can cause symptoms on kiwifruit, including *P. viridiflava, P. syringae* pv. *syringae* (Pss), and *P. syringae* pv. *actinidifoliorum*(Pfm; Visnovsky et al., 2019; González et al., 2024), yet little is known about how these might interact *in planta*. The Psa-kiwifruit pathosystem thus presents an opportunity to examine these interactions in the field across diverse, co-located host genotypes in *Actinidia* germplasm collections (Hemara et al., 2022; Hemara et al., 2024). Diverse *Actinidia* species have differing degrees of susceptibility/resistance to Psa3. While commercial *Actinidia chinensis* cultivars grown in monoculture are considered susceptible or tolerant, members of the *Leiocarpae* section of the *Actinidia* genus show resistance to Psa, with only select individuals from *A. arguta, A. macrosperma, A. valvata, A. polygama* and *A. melanandra* removed from New Zealand germplasm collections because of Psa infection (Datson et al., 2015). In particular, *A. arguta* and *A. melanandra* are known to recognise Psa through strong effector-triggered immune responses (Hemara et al., 2022; Hemara et al., 2024). While these species are not grown in large-scale commercial monocultures, they represent a wealth of genetic diversity in the *Actinidia* germplasm collections used for new cultivar development.

Following the establishment of Psa3 in New Zealand, a comprehensive genome biosurveillance programme was developed to monitor the evolution of Psa3 over time and to anticipate changes in virulence (Hemara et al., 2022; Hemara et al., 2024; Hemara et al., 2025). While the molecular basis for Psa3’s pathogenicity is well understood (McCann et al., 2013; Straub et al., 2018; Zhao et al., 2019; Jayaraman et al., 2020; Donati et al., 2020; Cellini et al., 2020; Hemara et al., 2022; Zhang et al., 2022; Jayaraman et al., 2023; Ishiga et al., 2023), little is known about how this pathogen may be interacting with other phyllosphere microbes, and even less about what constitutes the phyllosphere microbiome in *Actinidia* species outside of commercial monoculture. Sampling symptomatic plant material from *Actinidia* germplasm collections held at Plant & Food Research sites are a key part of tracking Psa’s evolution outside commercial orchard settings. In this work, this genome biosurveillance programme was leveraged to investigate more diverse *Pseudomonas* spp. isolated from the phyllosphere of symptomatic leaves from germplasm collections. In doing so, the following questions were addressed: What constitutes the bacterial community of diverse *Actinidia* species? What are the factors influencing community composition? Do the microbes associated with diverse *Actinidia* hosts cause the observed symptoms? Can the microbes resident on germplasm *Actinidia* form pathogenic consortia with Psa to increase disease? Or, alternatively, can these phyllosphere microbes form suppressive communities that reduce disease incidence or severity?

## Results

### *Pseudomonas syringae* dominates bacterial communities in symptomatic leaves of diverse *Actinidia* species

In December 2022, symptomatic leaves were sampled from diverse *Actinidia* species in germplasm collections at Plant & Food Research’s Kerikeri and Te Puke research orchards (Fig. 1A). Paired samples were taken, with leaf discs sampled from symptomatic lesions and asymptomatic areas of the same leaf (Fig. 1A). Despite widespread observation of leaves with angular, necrotic leaf spots, Psa was isolated from only 24 out of more than 200 symptomatic samples following selective isolation (Fig. 1B; Hemara et al., 2024). In spite of this low frequency of Psa isolation, when using DNA extracted directly from leaf homogenate, all symptomatic samples (bar four) tested positive for the presence of Psa by qPCR (Fig. 1C). In contrast, none of the asymptomatic samples tested positive (Fig. 1C). This also suggests that pathogen presence can be spatially limited across samples taken from the same leaf, mere centimetres apart (Fig. 1C). This low rate of Psa recovery was surprising, especially given the near-ubiquitous presence of Psa in the symptomatic samples. To investigate whether other bacteria could be contributing to symptom development, we sought to characterise the bacterial communities associated with disease symptomology across the sampled *Actinidia* germplasm.

**Figure 1.**
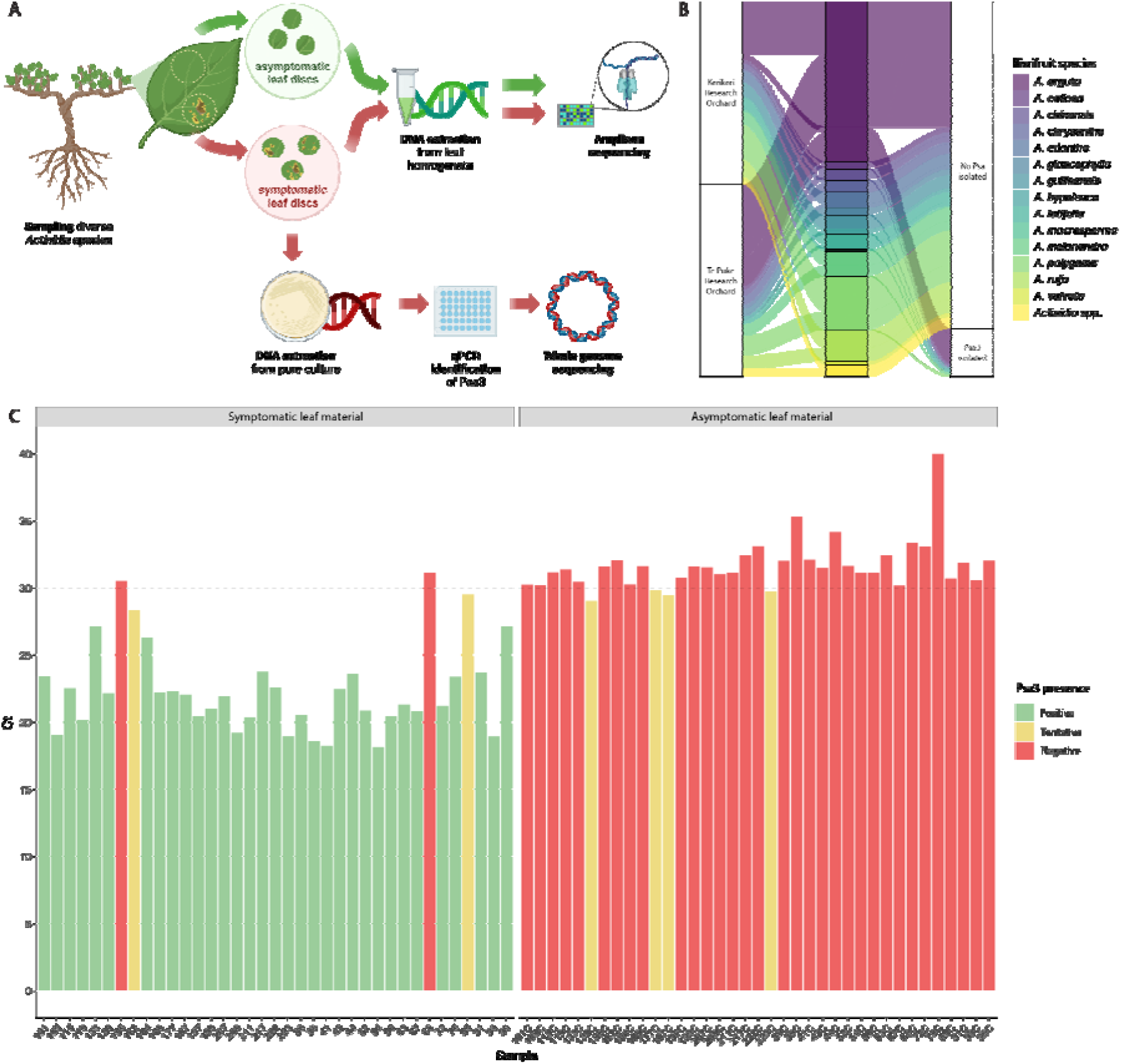
Low recovery of *Pseudomonas syringae* pv. *actinidiae* (Psa) isolates from symptomatic leaves in the *Actinidia* germplasm. (A) Germplasm sampling workflow schematic. Symptomatic leaf samples were taken from germplasm *Actinidia* vines. Symptomatic and asymptomatic leaf discs were sampled from each leaf and split into two sample sets. gDNA was extracted from leaf homogenate for quantitative PCR and amplicon sequencing. Leaf homogenate from symptomatic leaf discs was also plated on selective agar for *Pseudomonas* isolation. The resulting pure cultures underwent DNA extraction, qPCR testing for the presence of Psa, and whole genome sequencing. (B) Proportion of Psa isolations on selective agar from symptomatic leaf material, as confirmed by qPCR. (C) Presence of Psa in symptomatic leaf homogenate samples prepared for amplicon sequencing. Presence of Psa was assessed using qPCR with Psa F1/R2 primers (Andersen et al., 2017). Samples are grouped by sample status (symptomatic or asymptomatic). The horizontal dashed line indicates the cycle threshold (Ct) cut-off used to determine Psa presence.

Thirty-seven paired symptomatic and asymptomatic samples were selected from 16S amplicon sequencing, representing 13 *Actinidia* species that appeared symptomatic in the germplasm. V5-V9 primers demonstrated to be discriminatory against plant amplicons (Supplemtentary Fig. 1) were chosen to reduce off-target amplification of plastid DNA and to improve the effective sequencing depth (Mayer et al., 2021). The total number of sequencing reads per sample is reported, with most samples falling within an acceptable sequencing depth range of 75,000 to 150,000 reads (Supplementary Fig. 1; Caporaso et al., 2011). Amplicon sequencing controls showed the successful taxonomic classification of ONT-derived ASVs (Supplementary Fig. 2). All but six symptomatic samples were dominated by *Pseudomonas,* specifically amplicon sequence variants (ASVs) that appeared to fall within the Psa/Pth/Pfm clade in phylogroup 1 of *Pseudomonas syringae* (Fig. 2A). This is in line with qPCR reporting of Psa presence in virtually all symptomatic samples (Fig. 1C). These same ASVs (1,2,3,4,5,8,9, and 12) were present in the Psa3 V-13 gDNA control (Supplementary Fig. 2), suggesting that this was probably Psa3. These ASVs were also present in most asymptomatic samples, albeit at a much lower relative abundance than the corresponding symptomatic samples (Fig 2A). The Psa/Pth/Pfm ASVs 1 and 2 were some of the most abundant and appeared to be core members of the community across all samples (at a core threshold of 80%). Similarly, the Psa/Pth/Pfm ASVs 1,2,3,4,5 and 9 were core members of the symptomatic samples. The symptomatic status of samples, or the presence of this Psa/Pth/Pfm clade, resulted in reduced presence of other bacterial genera, as observed in compositional (Fig. 2A) and alpha-diversity plots (Supplementary Fig. 3). While symptomatic status and host species contribute to bacterial diversity, the contribution of region was statistically insignificant (Supplementary Figs 3, 4; Supplementary Table 1).

**Figure 2.**
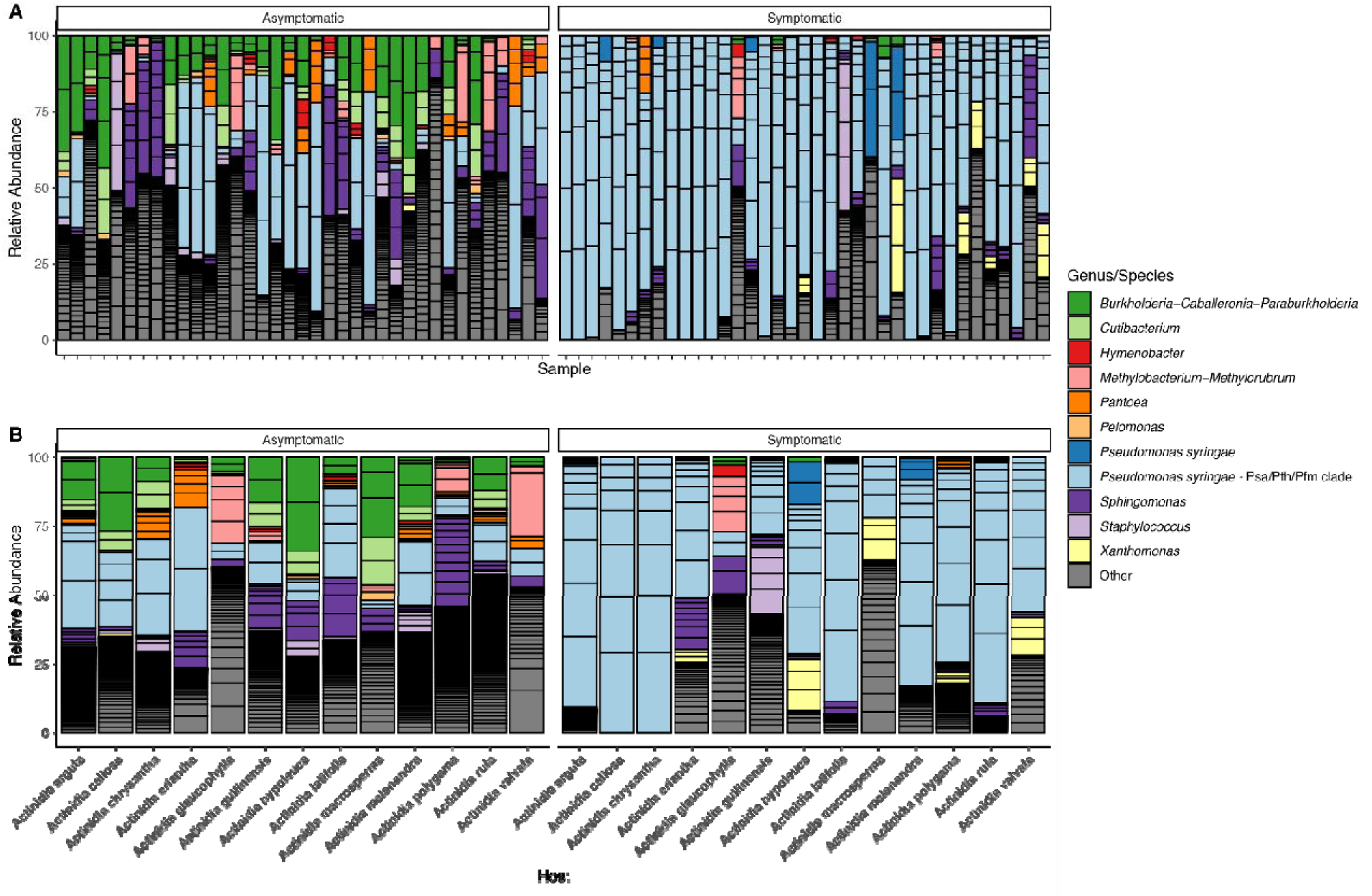
Dominant bacterial genera present in the *Actinidia* phyllosphere, by symptom status and host species. (A) Compositional bar plot of *Actinidia* phyllosphere communities by symptom status. The 50 most abundant amplicons sequence variants (ASVs) are represented, coloured by genus. (B) Compositional bar plot of *Actinidia* phyllosphere communities collated by *Actinidia* host. The 50 most abundant ASVs are represented, coloured by genus. ASVs identified as ‘*Pseudomonas syringae* – Psa/Pth/Pfm clade’ were identified by BLASTn. All other taxa are indicated by the group ‘Other’. Each black box within the bar indicates a different ASV.

While *Pseudomonas* dominated symptomatic samples, *Sphingomonas* and *Xanthomonas* ASVs were also present in some samples, with *Xanthomonas* present almost exclusively in symptomatic samples (Fig. 2A). Furthermore, the asymptomatic samples had diverse genera such as *Sphingomonas, Burkholderia, Cutibacterium, Methylobacterium,* and *Pantoea* (Fig. 2A). ASV7, assigned to *Burkholderia,* was a core member of the community of asymptomatic samples. Other ASVs of note that appeared to be differentially abundant in asymptomatic tissue is *Cutibacterium acnes* (ASVs 13 & 14). Sequencing of our mock community and no-template control (NTC) indicated that this was not contamination during the extraction and sequencing processes (Supplementary Fig. 2).

When samples were grouped by host, it was apparent that some species had distinct bacterial communities (Fig. 2B). For example, *A. callosa* and *A. chrysantha* symptomatic samples were dominated by the Psa/Pth/Pfm clade, while the corresponding asymptomatic samples were diverse (Fig. 2B). Similar observations can be made for *A. rufa* and *A. latifolia,* albeit to a lesser extent (Fig. 2B). *A. glaucophylla* and *A. guilinensis* show the highest alpha-diversity values (Supplementary Fig. 4), reflected in high genera diversity and low abundance of Psa/Pth/Pfm (Fig. 2B). Interestingly, the bacterial communities of symptomatic *A. glaucophylla* and *A. guilinensis* samples were similar to those of asymptomatic samples from other hosts, with lower abundance of *Pseudomonas* (Fig. 2B). Notably, *A. guilinensis* symptomatic samples had abundant *Staphylococcus* ASVs (Fig. 2B). Finally, symptomatic *A. glaucophylla, A. guilinensis,* and, to a lesser extent, *A. eriantha, A. hypoleuca,* and *A. macrosperma,* had relatively less Psa/Pfm/Pth than other species (Fig. 2B). Interestingly, hosts on which we have seen Psa3 effector loss variants emerge, including *A. arguta* and *A. melanandra,* had high abundance of Psa/Pth/Pfm despite Psa recognition (Hemara et al., 2022; Hemara et al., 2024). This could be explained if resistance-escaping, effector loss variants have spread in the germplasm, thereby contributing to this relatively high abundance. However, only sample 39 from *Actinidia melanandra* yielded an effector loss variant – Psa3 X_477, which has lost *avrRpm1a* (Hemara et al., 2024). The limited isolation of effector loss variants from this sample suggests that the majority of these Psa/Pth/Pfm ASVs are Psa3 lineages carrying the full ETI-eliciting effector repertoire.

### Diverse *Pseudomonas* spp. isolated from symptomatic material carry genes belying pathogenic potential

While ASV analysis has revealed dominant genera and, in some instances, species and specific lineages, partial 16S amplicons do not allow full taxonomic resolution to the pathovar or biovar level. Diverse *Pseudomonas* were isolated from symptomatic plant material (Fig. 1A, 1B), despite the dominance of the Psa/Pth/Pfm clade in leaf homogenate samples. The isolation strategy used in this work indiscriminately captured both epiphytic and endophytic bacteria from symptomatic tissue, making it plausible that some of these isolates could be colonising the apoplastic space and contributing to disease. Whole genome sequencing (WGS) was performed on a subset of these isolates, allowing us to assign more detailed taxonomy through comparison to a core gene alignment of *P. syringae* and plant-associated *Pseudomonas* reference genomes (Supplementary Fig. 5). Of the 16 germplasm isolates analysed, 12 were categorised within the *P. syringae* species complex: five isolates fell into phylogroup 3, three into phylogroup 2, and one each into phylogroups 7, 11, and 13 (Supplementary Fig. 5). One further isolate—HP_008—remained unassigned to a phylogroup within the *P. syringae* species complex, clustering with PsyCFBP13 (Supplementary Fig. 5). HP_009 fell within the *P. protegens* subgroup, HP_015 within the *P. fluorescens* subgroup, and HP_001 and HP_012 within the *P. lutea* group (Supplementary Fig. 5). Interestingly, when the 16S sequences of these germplasm isolates were compared to our *Pseudomonas* ASVs, these isolates universally appeared to be present only in one or two samples, and at low abundance.

To ascertain the pathogenic potential of these HP germplasm isolates, the isolates’ genomes were firstly screened for key virulence factors, including pathogenicity-associated islands (PAIs) encoding type III secretion systems for effector delivery, type III secreted effector repertoires, and pathogen-associated secondary metabolite biosynthetic pathways (Xin et al., 2018). As expected, all seven isolates from canonical *P. syringae* phylogroups had a T3SS R-PAI (Fig. 3). In the non-canonical *P. syringae* phylogroups, two further isolates had S-PAIs, and HP_014 carried a PG13 A-PAI (Fig. 3). Outside of the *P. syringae* species complex, some isolates carried T-PAIs (Fig. 3). Overall, only three isolates had no identifiable PAI (Fig. 3). Twenty-five families of effectors were found in the genomes of germplasm isolates (Fig. 3). The most frequent effector family was HopB2, followed by AvrE1. However, across all these genomes, as well as the manually curated Psa3 V-13 assembly, HopB2 lacked a HrpL box. Prior to effector family reassignment (Dillon, Almeida, et al., 2019), HopB2c was previously annotated in the Psa3 V-13 genome as Aconitase A, a core gene in the lipopolysaccharide locus, and may be mischaracterised as an effector. All isolates from canonical *P. syringae* phylogroups carried AvrE1, HopM, HopAA1, and HopI1 (Fig. 3), which have previously been identified as core effectors (Lindeberg et al., 2012; Dillon, Almeida, et al., 2019). While many isolates carried effectors from the same family, these effectors often belonged to different subtypes (Supplementary Table 2). HP_006 and HP_011, both from phylogroup 2, carried the highest number of effectors at thirteen (Fig. 3). Conversely, non-canonical *P. syringae* HP_014 carried only a single effector (Fig. 3). Non-canonical HP_008 and isolates outside the *P. syringae* species complex did not carry any *P. syringae* -related effectors, despite HP_012 and HP_001 appearing to carry a canonical and alternate T-PAI, respectively (Fig. 3). This potentially indicates a limitation in the ability to detect effectors by homology with distantly related strains (Fig. 3).

**Figure 3.**
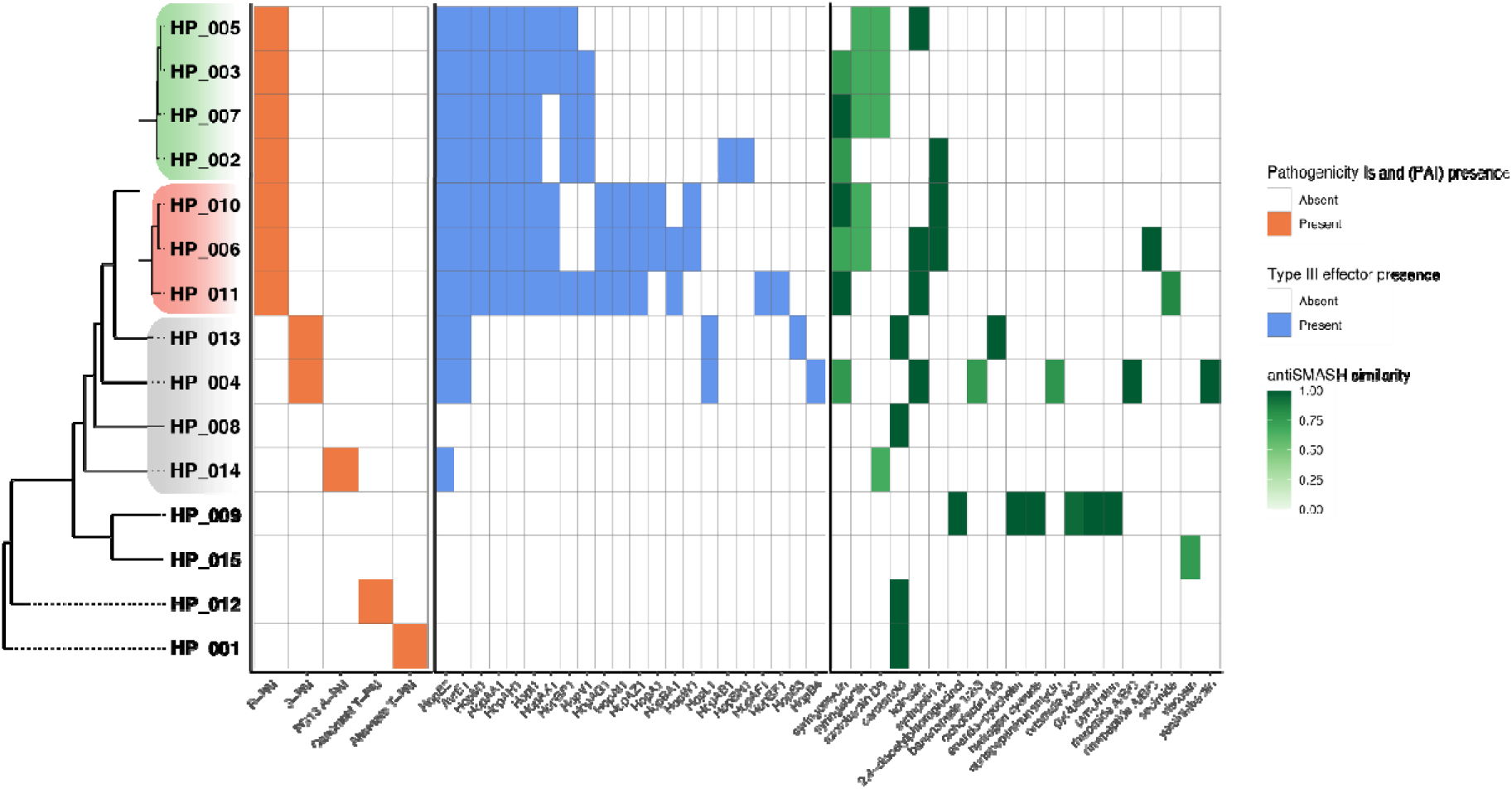
Pathogenicity-associated traits found across germplasm *Pseudomonas* isolates. Phylogeny colours represent *Pseudomonas syringae* phylogroups 2 (red) and 3 (green), as well as non-canonical *P. syringae* (grey). Pathogenicity island types expected in the *Pseudomonas syringae* species complex. *P. syringae* effectors, as defined in Dillon et al. (2019), are ordered by prevalence. Gene clusters with ≥60% similarity to known secondary metabolites, as identified by antiSMASH (version 7.0; Blin et al., 2023).

The presence of secondary metabolite biosynthetic gene clusters was also assessed using antiSMASH (Blin et al., 2023). The most common secondary metabolite type across all isolates was non-ribosomal peptide synthetases (NRPSs). From these NRPSs, there were at least twelve biosynthetic gene clusters of interest found across these germplasm isolates spanning a variety of known toxins (anti-host and anti-microbial), siderophores, and surfactants (Fig. 3).

### Host species influence interactions between *Pseudomonas* isolates and Psa3

Non-Psa *Pseudomonas* spp. appear to be minor components of the phyllosphere community yet possess virulence components that could be contributing to the increased pathogenicity of Psa in Psa-resistant germplasm. To examine the pathogenic potential of these germplasm-derived *Pseudomonas* spp., isolates were assayed for *in planta* growth across diverse kiwifruit hosts, including *A. arguta* AA07_03, *A. melanandra* ME02_01, and *A. polygama* PC02_01, as well as the susceptible kiwifruit model plant *A. chinensis* var. *chinensis* ‘Hort16A’. ‘Hort16A’ was used to compare plate count and qPCR quantification validating the use of isolate-specific primers for relative quantification of *in planta* growth 7 dpi (Supplementary Fig. 6). Interestingly, HP_013 from phylogroup 7 appeared to grow as well as virulent Psa3 V-13 on this susceptible host (Supplementary Fig. 6). Isolates HP_004, HP_014, HP_001, and HP_007 had comparable growth to the non-pathogenic epiphyte Pfm, infecting ‘Hort16A’ poorly (Supplementary Fig. 6). The remaining three isolates tested (HP_015, HP_009, and HP_006) showed intermediate infection of ‘Hort16A’. (Supplementary Fig. 6).

On the germplasm hosts *A. arguta, A. melanandra,* and *A. polygama,* isolates appeared to have relatively lower bacterial growth than on susceptible ‘Hort16A’ (Fig. 4A, 4B, 4C; Supplementary Fig. 6). The majority of these isolates were non-pathogenic in single-isolate infections, as defined by pathogenicity as the qualitative ability to infect and cause symptoms on a host (Sacristan & Garcia-Arenal, 2008). Furthermore, it appears that isolates derived from any given host do not have any ‘home advantage’ (Fig 4A, 4B, 4C). On *A. arguta,* only a few isolates had significantly different growth from that of recognised, and thus avirulent, Psa3 V-13 (Fig. 4A). HP_004 was significantly different from Psa3 V-13 with slightly higher *in planta* growth, while HP_014 grew the least by a large margin (Fig. 4A). On *A. melanandra,* germplasm isolates were more similar to one another: isolates HP_006, HP_001, and HP_015 had reduced growth compared with the recognised Psa3 V-13, with growth of all other isolates not significantly different from this control (Fig. 4B). Finally, on *A. polygama,* all isolates demonstrated relatively low growth, and most were not significantly different from one another (Fig. 4C). Across all germplasm hosts, none of the isolates, or the controls, showed pathogen-associated levels of growth (Fig. 4A, 4B, 4C). Curiously, however, a distinct leaf collapse and tissue necrosis phenotype was observed when ‘Hort16A’, *A. arguta,* and *A. melanandra,* but not *A. polygama,* were infected with HP_013, (Supplementary Fig. 9).

**Figure 4.**
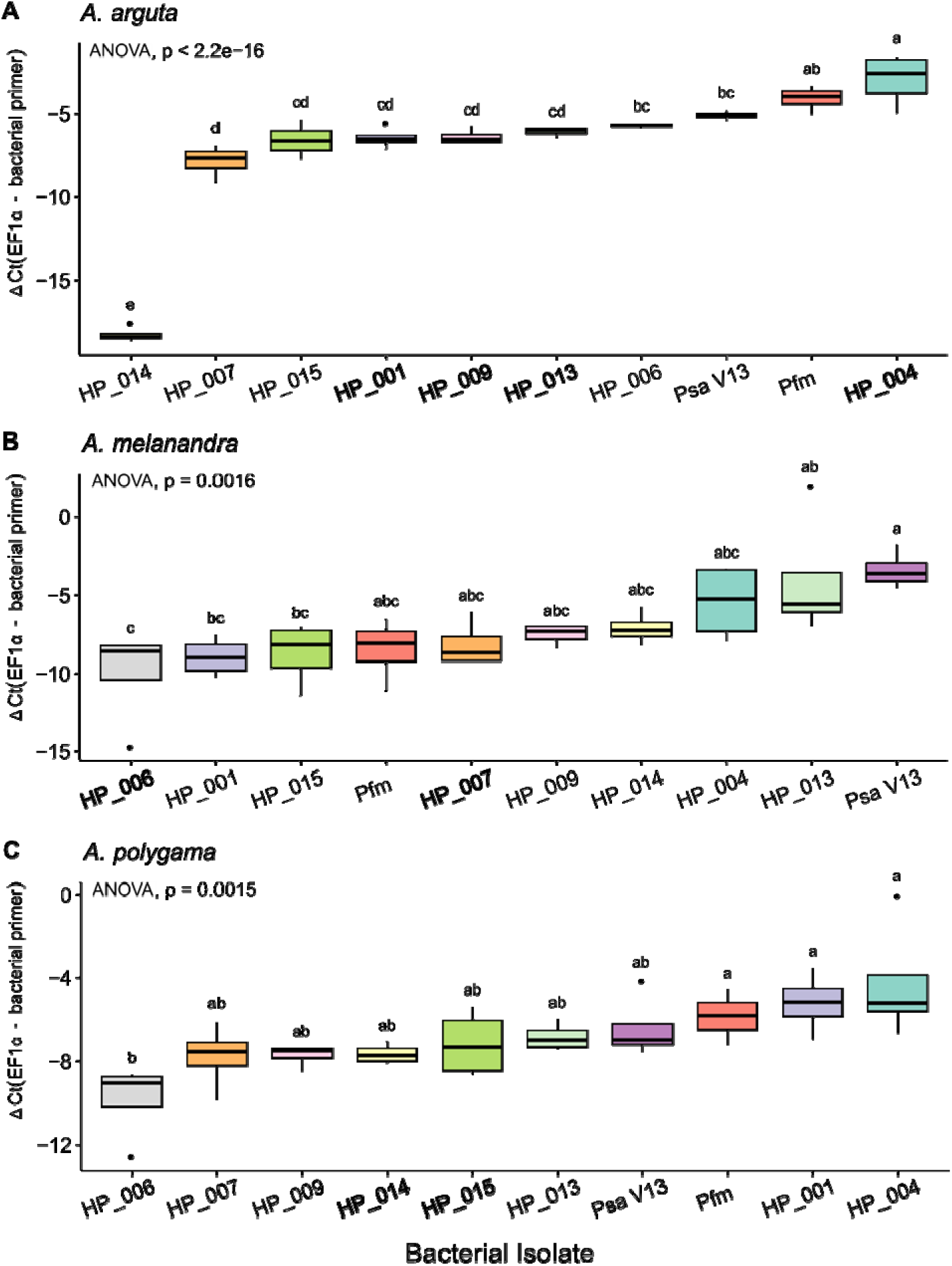
Results of qPCR analysis of bacterial growth of germplasm isolates in three species o f *Actinidia* plantlets at 7 dpi. Plantlets were flooded with 10^6^ CFU/mL of each bacterial isolate. Bacterial growth was quantified by quantitative real-time PCR using isolate-specific primers, and normalising the Ct to a plant housekeeping gene, EF1a to get a ΔCt value. The isolates were tested against A) *Actinidia arguta,* B) *Actinidia melanandra,* and C) *Actinidia polygama* plantlets. The isolates highlighted with bold font on the x-axis were isolated from that respective species. Black bars represent the median values, box whiskers indicate total range of four technical replicates. Compact letter display above each box indicates significant differences between bacterial isolates using Tukey’s Honestly Significant Difference (HSD) post hoc at 5%.

These germplasm isolates remain curious; while not particularly abundant in amplicon sequencing data, these *Pseudomonas* spp. were isolated from symptomatic material from which we expected to recover Psa. While they have pathogenic potential, as suggested by their genomic characteristics, the vast majority did not have increased *in planta* growth over neither the avirulent Psa3 V-13 or epiphytic Pfm on resistant germplasm species. The lack of independent colonisation by these isolates raised the possibility that they may only be opportunistically present in detectable/recoverable quantities during Psa-mediated infection, forming a commensal syndemic interaction (hypothesis 1). Alternatively, because of the low incidence of Psa-infection *per se* in these genotypes, these strains may be facilitating the growth of Psa through formation of mutualistic syndemic infections as suspected in previous studies (hypothesis 2) (Purahong et al., 2018; Sadhukan et al., 2024). To test the potential for either of these dynamics between these germplasm-derived non-Psa isolates and Psa, 1:1 ratios of co-inoculation of each of these isolates and Psa were set up on the *Actinidia* species from which these non-Psa isolates were derived. On *A. melanandra,* co-inoculation did not have a strong effect on the growth of either the germplasm isolates or Psa3 V-13 (Fig. 5). Compared with individual inoculation, there was no difference in growth when HP_006 and Psa3 V-13 were co-inoculated (Fig. 5A). HP_007 decreased the growth of Psa3 V-13 when co-inoculated (*p* = 0.0022), but its own growth was unaffected when co-inoculated (Fig. 5B). Similar dynamics were observed when Psa3 V-13 was co-inoculated with the control Pfm (*p* = 0.033; Fig. 5C). Curiously, the inverse was observed when isolates HP_014 and HP_015 were tested on *A. polygama* (Fig. 6). On *A. polygama,* the co-inoculation of HP_014 and Psa3 V-13 resulted in the increased growth of both isolates compared with individual inoculation, suggesting syndemic growth of Psa and HP_014 and evidence for hypothesis 2 (*p* = 0.00051 and *p* = 0.01, respectively; Fig. 6A). When co-inoculated with HP_015, the growth of Psa3 V-13 also increased (*p* = 0.039), although HP_015 remained unaffected again suggesting evidence for hypothesis 2 (Fig. 6B). Neither Pfm nor Psa3 V-13 were affected when co-inoculated into *A. polygama,* suggesting that host genotype may influence this syndemic interaction outcome (Fig. 6C; Supplementary Table 3).

**Figure 5.**
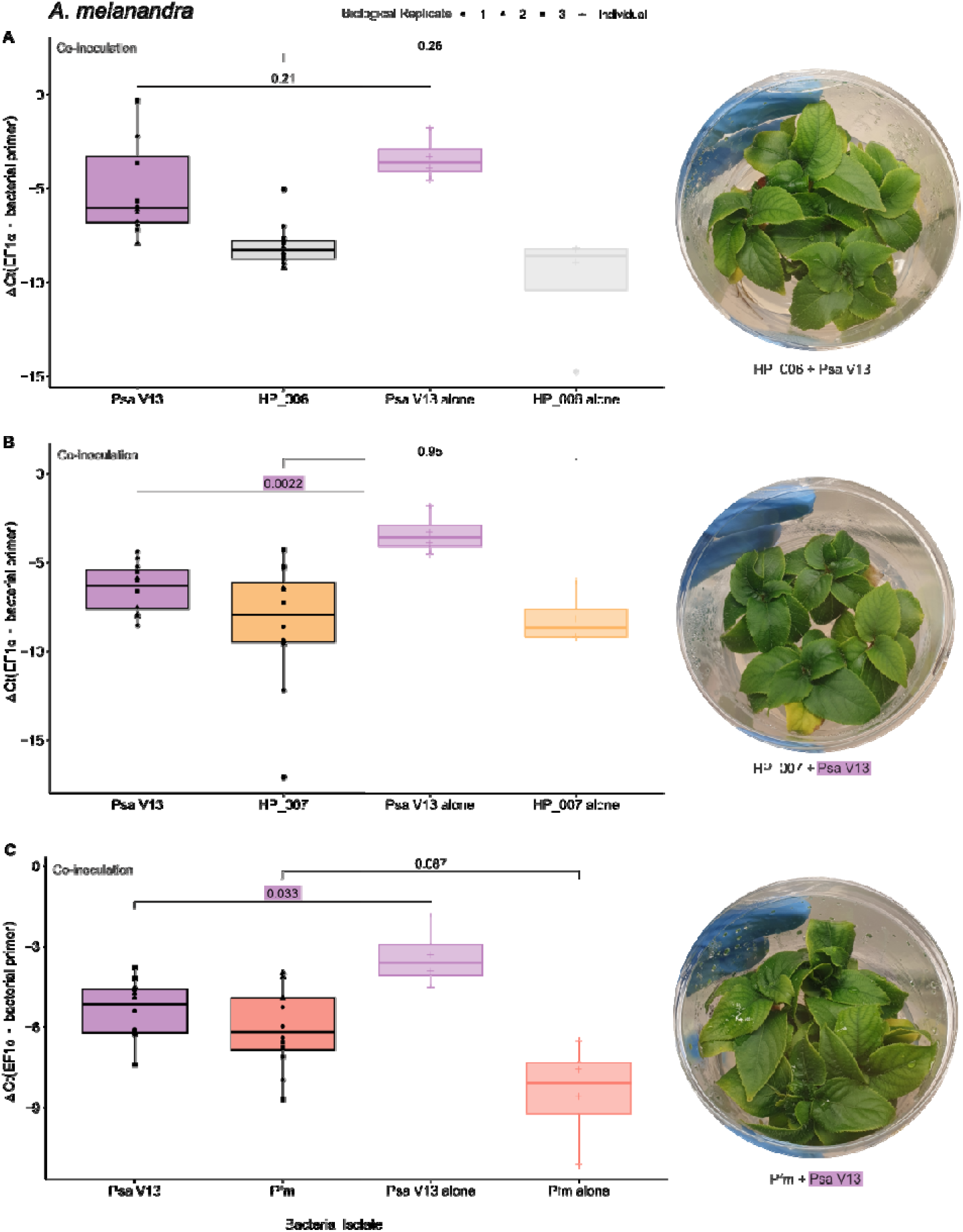
Results of qPCR analysis of virulence of germplasm isolates with *Pseudomonas syringae* pv. *actinidiae* (Psa) on *Actinidia melanandra* at 7 dpi. Plantlets were flooded with 10^6^ CFU/mL of bacterial isolates together. Bacterial growth was quantified by quantitative real-time PCR using isolate-specific primers, and normalising the Ct to a plant housekeeping gene, EF1a to get a ΔCt value. A) HP_006 and Psa3 V-13 (*p* -value generated with Wilcoxon’s test), B) HP_007 and Psa3 V-13 (*p* -value generated with Wilcoxon’s test), and C) *Pseudomonas syringae* pv. *actinidifoliorum* (Pfm) and Psa3 V-13 (*p* -value generated with Welch’s t-test). Each pane compares the results of the *in plant a* competition assay with each isolate in the same host alone using a pairwise significance test. Representative plantlet images at 7 dpi are displayed next to the relevant assay. Significant results are highlighted with the colour of the variable to which it pertains.

**Figure 6.**
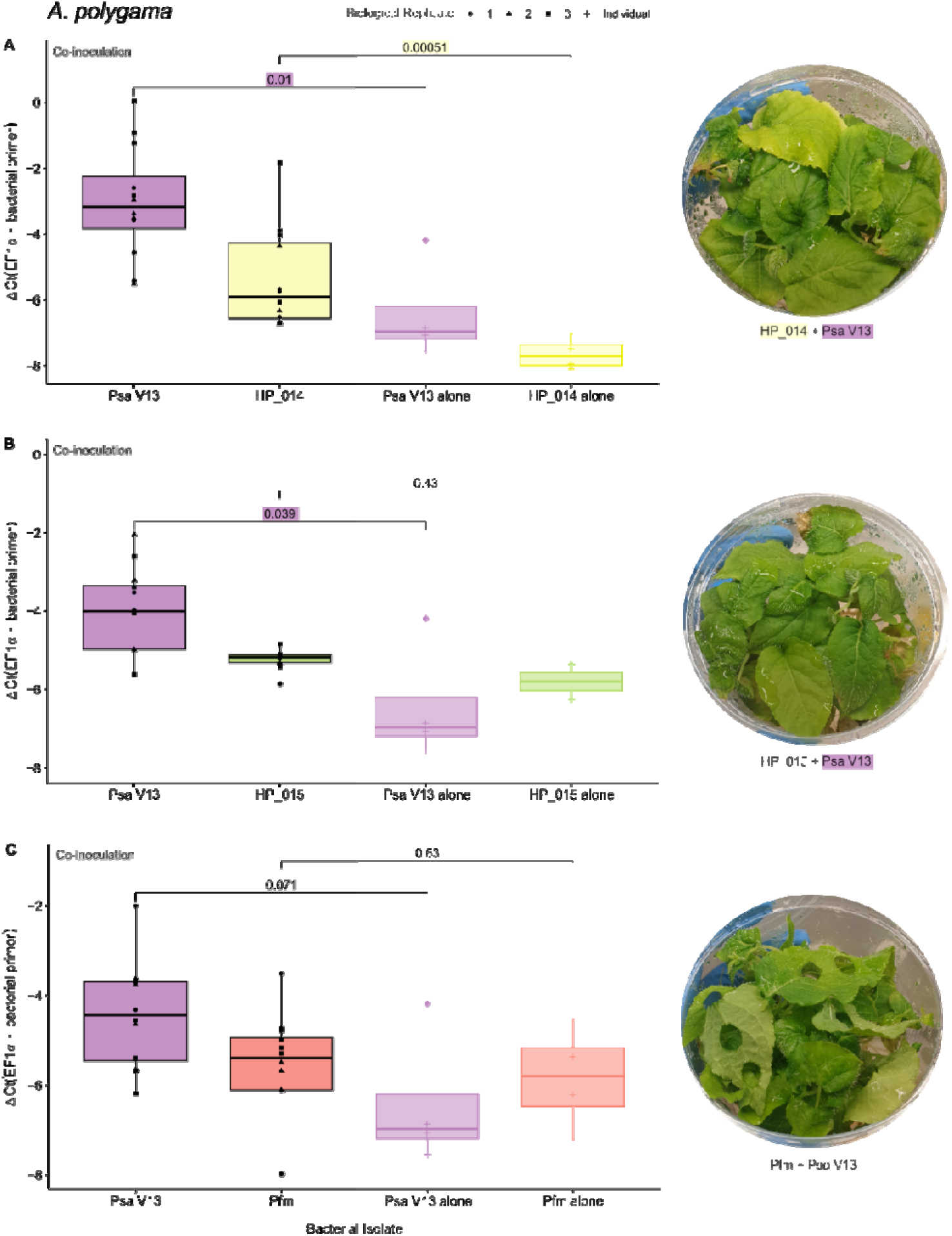
Results of qPCR analysis of virulence of germplasm isolates in competition wit h *Pseudomonas syringae* pv. *actinidiae* (Psa) on *Actinidia polygama* at 7 dpi. Plantlets were flooded with 10^6^ CFU/mL of each bacterial isolate. Bacterial growth was quantified by quantitative real-time PCR using isolate-specific primers, and normalising the Ct to a plant housekeeping gene, EF1a to get a ΔCt value. A) HP-014 and Psa3 V-13, B) HP_015 and Psa3 V-13, and C) *Pseudomonas syringae* pv. *actinidifoliorum* (Pfm) and Psa3 V-13. Each pane compares the results of the *in planta* co-inoculation assay with each isolate in the same host alone, with *p*-values generated using Welch’s t-test. Representative plantlet images at 7 dpi are displayed next to the relevant assay. Significant results are highlighted with the colour of the variable to which it pertains.

Notably, four isolates were derived from *A. arguta,* and co-infections revealed several to have increased growth when co-inoculated with Psa3 V-13 (Fig. 7). Interestingly, *A. arguta* was the only host to exhibit disease symptoms across any of the co-inoculation experiments, with HP_004 and HP_013 co-inoculations greatly increasing the *in planta* growth of both Psa3 V-13 and the germplasm isolate, also supporting hypothesis 2 (Fig. 7B, 7D). Curiously, this same increase was also observed for the Pfm control (Fig. 7E), which was not observed on *A. melanandra* (Fig. 5C) or *A. polygama,* again suggesting host-specificity of syndemic associations (Fig. 6C). Only co-inoculation with HP_001 saw no change for either isolate during co-inoculation (Fig. 7A), while HP_009 slightly decreased the growth of Psa3 V-13 when co-inoculated (Fig. 7C). Fittingly, these plants appeared asymptomatic following co-inoculation.

**Figure 7.**
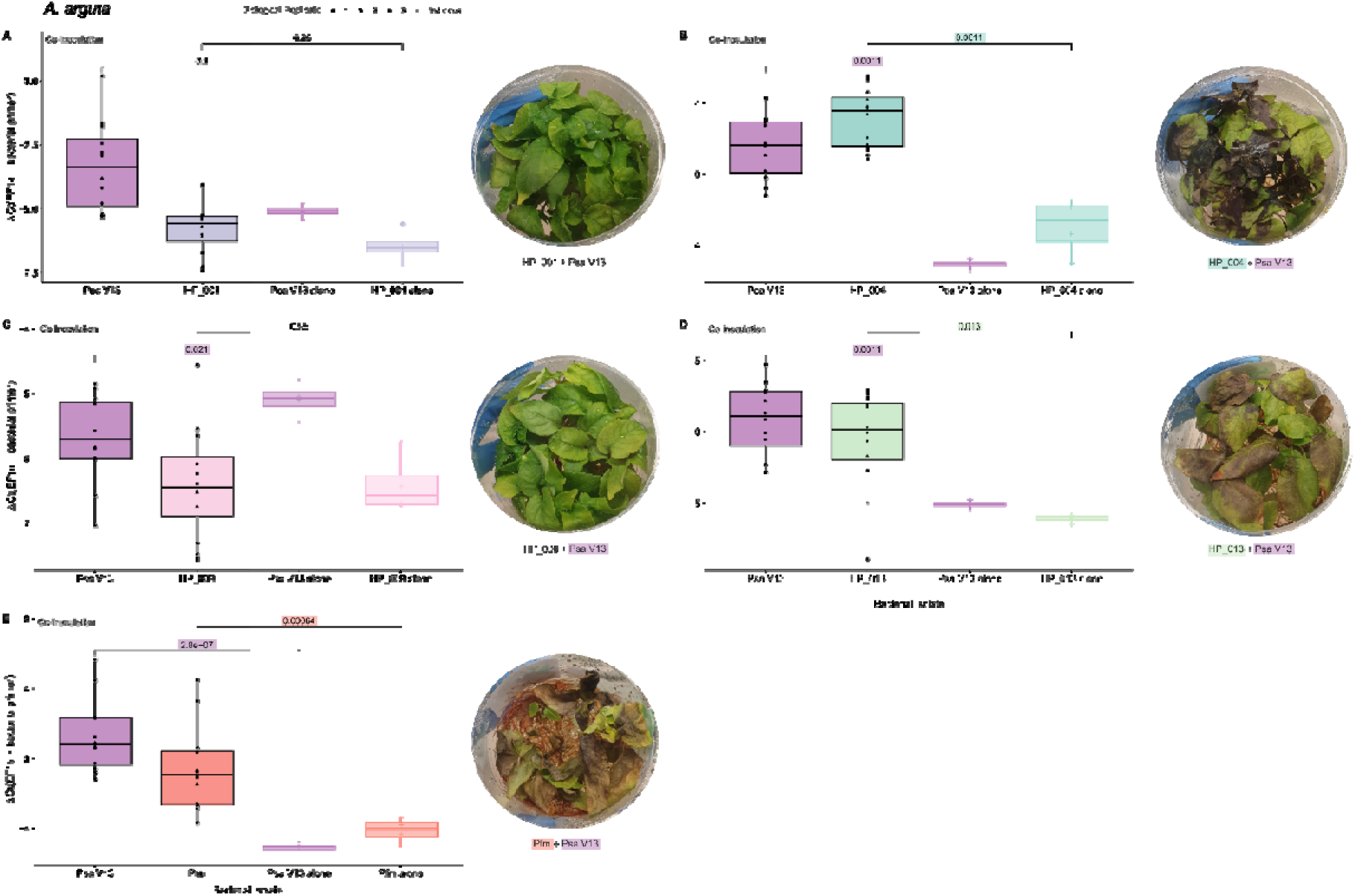
Results of qPCR analysis of virulence of germplasm isolates in competition wit h *Pseudomonas syringae* pv. *actinidiae* (Psa) on *Actindia arguta* at 7 dpi. Plantlets were flooded with 10^6^ CFU/mL of each bacterial isolate. Bacterial growth was quantified by quantitative real-time PCR using isolate-specific primers, and normalising the Ct to a plant housekeeping gene, EF1a to get a ΔCt value. A) HP_001, B) HP_004, C) HP_009, D) HP_013, and E) *Pseudomonas syringae* pv. *actinidifoliorum* (Pfm). Each pane compares the results of the *in planta* co-inoculation assay with each isolate in the same host alone, with *p*-values generated using Welch’s t-test. Representative plantlet images at 7 dpi are displayed next to the relevant assay. Significant results are highlighted with the colour of the variable to which it pertains.

## Discussion

This work addressed the need to understand the basis of disease symptomology arising on Psa-resistant *Actinidia* germplasm, given the low recovery of Psa isolates. While Psa does grow more slowly than some other *P. syringae* lineages on agar, taking 30–48 hours for colonies to appear following room temperature incubation (Tyson et al., 2018), the lack of many resistance-breaking Psa3 isolates suggested this could not necessarily explain all the symptomatic material (Hemara et al., 2024). To this end, we characterised the associated bacterial phyllosphere communities. Unsurprisingly, ASV characterisation demonstrated that Psa infection affects the composition and diversity of bacterial communities. Microbiome changes in disease settings appear to be a universal phenomenon (Agler et al., 2016; Berendsen et al., 2018; Gao et al., 2021; Díaz-Cruz & Cassone, 2022; Li et al., 2022), and is also consistent with previous observations in the kiwifruit phyllosphere (Purahong et al., 2018; Ares et al., 2021). Symptomatic samples were dominated by *P. syringae* from the Psa/Pth/Pfm clade, which we assume to represent Psa3. The *Pseudomonas* genus, when present, was almost entirely dominated by this *P. syringae* phylogroup 1 clade. Interestingly, there was little detection by metabarcoding of other *Pseudomonas* spp. known to be present in New Zealand kiwifruit orchards, such as *P. syringae* phylogroup 3a isolates which have been found epiphytically on kiwifruit (Straub et al., 2018; Visnovsky et al., 2019). The same holds true for *Pss* and *P. viridiflava,* which were previously found in *A. chinensis* var. *chinensis* and *A. chinensis* var. *deliciosa* infected with Psa (Purahong et al., 2018). Beyond *Pseudomonas, Xanthomonas* ASVs were present almost exclusively in symptomatic samples. Xanthomonads are known to opportunistically form pathogen complexes with *Pseudomonas* in disease settings, such as with *P. syringae* in pepper and tomato infections (Gitaitis et al., 1987). The near-exclusive occurrence of *Xanthomonas* in symptomatic samples could suggest that they have a role in disease or take advantage of Psa presence to opportunistically co-colonise leaf tissue. Overall, the ability of pathogens to manipulate their host ultimately affects microbial competition and resource availability, affecting microbial diversity likely through both direct resource utilisation and effector delivery to modify the host’s metabolome (Cai et al., 2023; Schlechter et al., 2023).

The most diverse communities were observed in asymptomatic leaf tissue, with infection status the most significant factor reducing diversity in community composition in this study. While the Psa/Pth/Pfm lineage was often present in these samples (possibly as latent/emerging infections), asymptomatic samples were also dominated by *Cutibacterium, Sphingomonas, Burkholderia, Methylobacterium,* and *Pantoea*. *Cutibacterium acnes* has been found in association with grapevines as an endophyte, where it appears to have diverged from human-associated strains 7,000 years ago when grape was domesticated (Campisano et al., 2014). In *P. syringae* - infected tobacco, treatment with biological control agents (BCAs) resulted in greater presence of *Sphingomonas* and *Methylobacterium,* which are suggested to have plant-protective traits through competition and defence priming, respectively (Yang et al., 2024). *Pantoea* have been used in biocontrol against *P. syringae,* as have other *Pseudomonas* spp. (Yang et al., 2024). Recent work has demonstrated that the early recruitment of eubiotic microbiota aids the development of plant immune responses against pathogens in *Arabidopsis,* without which the plant becomes susceptible to disease (Paasch et al., 2023). However, pathogen perturbation of the wider microbiome, through mechanisms such as type VI secretion system-mediated competition or antimicrobial activity, can lead to dysbiosis and disease (Chen et al., 2020; Smith et al., 2020). Conversely, the phyllosphere microbiome can also change to benefit the plant after pathogen invasion, reducing dysbiosis and disease severity (Ehau-Taumaunu & Hockett, 2023). These plant-protective bacteria may act by inducing host resistance to inhibit pathogen growth and disease (Bashir et al., 2022). Further still, our focus on the potential contribution of these phyllosphere communities to pathogen resistance also overlooks the other beneficial contributions they make, including phytohormone production, nitrogen fixation, and the solubilisation of phosphate and potassium (Bashir et al., 2022). The role of phyllosphere bacteria thus cannot be downplayed – they represent a large source of functional diversity from which plant hosts may gain benefits, including pathogen resistance. That these different communities exist mere centimetres apart shows the influence that pathogen presence has on the community at a highly local level. Overall, these results suggest a highly localised community on the leaf surface, with *Pseudomonas* spp. dominating symptomatic tissue while more diverse communities occupy other, asymptomatic portions of the leaf.

Plant genotype is another important contributor to microbiome composition (Wagner et al., 2020; Bashir et al., 2022). In this work, host genotype was found to be a significant factor, in line with previous studies in commercial monoculture settings that have shown different varieties to have significantly different microbiome compositions (Purahong et al., 2018; Louisson et al., 2023). *A. chinensis* var. *chinensis* exhibited higher diversity than *A. chinensis* var. *deliciosa* (Purahong et al., 2018). Notably, *Pseudomonas* spp. were thirteen times more abundant on var. *deliciosa* than on var. *chinensis* (Purahong et al., 2018). Further still, differences in phyllosphere microbiomes of these two kiwifruit varieties were more pronounced when a pathogen was present (Purahong et al., 2018). Pathogen-infected *A. chinensis* var. *deliciosa* microbiomes were dominated by over 75% *Pseudomonas* spp., compared with less than 25% in infected *A. chinensis* var. *chinensis* (Purahong et al., 2018). This suggests that pathogen presence affects the phyllosphere microbiome in a host-dependent manner, which supports similar observations in this work. Differences in the microbiome were attributed to trichome distribution and structure between the two species (Purahong et al., 2018). Based on the phenotypic differences observed in leaves from diverse *Actinidia* spp., it is plausible that a similar phenomenon could be at play in the germplasm. In the tomato phyllosphere, trichomes form genotype-specific microbial hotspots, boosting bacterial diversity when present, particularly *Sphingomonadacea* and *Burkholderiaceae* (Kusstatscher et al., 2020). Furthermore, it has recently been demonstrated that *Erwinia amylovora* enters apple leaf tissue through natural wounds produced by trichome abscission, which provides another possible explanation as to how trichomes may influence phyllosphere communities (Millett et al., 2024). Several factors affecting the plant-associated microbiota remain to be explored for diverse genotypes in *Actinidia*. Seasonality is a trait that in conjunction with host genetics determines microbiome composition (Ares et al., 2021; Howe et al., 2023). Future work could also look at the effects of plant genetics from the lens of geographical distribution with shared geography affecting colonisation by microbial community members when overlaid on host genetics (Busby et al., 2013). Taken together, these works highlight the important influence plant architecture and micro-niches, plant genetics, co-local geographic distribution of hosts, and seasonality may have on community composition (Kusstatscher et al., 2020; Howe et al., 2023; Busby et al., 2013).

Notably, A. glaucophylla and A. guilinensis were not dominated by *Pseudomonas* spp., with diverse communities observed across both symptomatic and asymptomatic samples. It could be that these species have hyper-resistance against Psa, thus suppressing *Pseudomonas* abundance as observed at the genus level. Another hypothesis could be that, rather than relying on R gene-mediated effector recognition, these species modulate their microbiomes using ‘M’ genes (Zhan & Wang, 2024). ‘M’ genes are able to recruit certain genera and maintain microbiome homeostasis, as demonstrated in rice where 4-hydroxycinnamic acid enriches for *Pseudomonas* spp. (Su et al., 2024). Although this ‘M’ gene concept has yet to be demonstrated in other species, it presents an interesting possibility for investigation. Interestingly, *A. arguta* has recently been demonstrated to produce hydroxycinnamic acids (Kovalska et al., 2023; Laima Česonienė et al., 2024). Exploring the contribution of both R genes to disease resistance and potential ‘M’ genes to disease suppression is a critical avenue of future research for kiwifruit breeding pipelines seeking to tap into the genetic potential held within the germplasm. Completely asymptomatic germplasm vines, which were not sampled in this work, could also be a valuable source of protective bacterial strains. Further still, further genome sequencing and mining of the *Actinidia* germplasm may reveal whether ‘M’ genes are indeed present, and whether selective microbiome recruitment could be a viable strategy to suppress disease in the *Actinidia* phyllosphere. Alternatively, differences in *Pseudomonas* abundance across hosts could be attributed to the specific variants present. While we propose that some species may have little or no presence of the Psa/Pth/Pfm clade owing to hyper-resistance, we do not observe this same phenomenon in *Actinidia* species we know to have strong effector-triggered immunity against Psa, such as *A. arguta* and *A. melanandra* (Hemara et al., 2022, 2024). Further still, these studies have highlighted the emergence of effector loss isolates that partially escape host immunity and have improved *in planta* growth (Hemara et al., 2022, 2024). Whether the Psa population present on particular hosts represents ‘wild-type’ Psa3 or effector loss variants could further explain why this Psa/Pth/Pfm clade is dominant on some hosts and not others. However, isolation efforts suggest that effector loss variants were not widespread within this sample set (Hemara et al., 2024). Other factors are likely at play as well – for example, phyllosphere bacterial communities in *A*. *chinensis* var. *chinensis* and *A. chinensis* var. *deliciosa* were also influenced by stochastic assembly processes, particularly drift and dispersal (Louisson et al., 2023). Increasing the sample size of each species and taking advantage of duplicate accessions planted out in germplasm collections at different locations (New Zealand sites in Kerikeri, Te Puke, and Motueka) will allow further studies to better understand the contributions of host genotype, pathogen presence, and stochastic assembly to microbiome composition.

Whole genome sequencing of *Pseudomonas* isolates revealed the presence of potential pathogen-associated genomic features, including well characterised T3SS effectors and several secondary metabolites with potential toxin functions. The effector repertoires of germplasm isolates were generally small compared wit those of Psa strains which have an average of 39 (McCann et al., 2013), and Pfm ICMP 18803 which carries 31 (Templeton et al., 2023). This could suggest that the isolates are epiphytic and non-pathogenic, or that they may contribute to symptomology through other means, such as toxins. The widespread presence of AvrE1 suggests a common basic ability among these isolates from symptomatic tissues to induce apoplastic wetting (Nomura et al., 2023). Some germplasm isolates carried few effectors, such as the *P. viridiflava* isolate HP_013 which encodes only AvrE1. Accordingly, these isolates may not be able to gain entry to the apoplast or suppress plant immunity alone, thus making a partnership with Psa3 V-13 beneficial. However, Psa3 V-13 is recognised on these *Actinidia* species (Hemara et al., 2022, 2024), implying that isolates like HP_013 must also be providing colonisation or immune suppression advantages, to improve *in planta* growth outcomes for both isolates when co-inoculated.

Several of the secondary metabolite biosynthesis pathways identified in these genomes have the potential to aid plant infection. Syringomycin and syringafactin were found in a number of canonical *P. syringae* isolates. Syringomycin contributes to virulence in *P. syringae* by forming lethal pores in the lipid bilayers of host cells (Bender et al., 1999; Scholz-Schroeder et al., 2001). Syringafactin is proposed to increase permeability of the leaf cuticle for nutrient diffusion and possibly facilitating the action of other phytotoxins (Girard et al., 2020). Syringolin A, found in isolates HP_002, HP_006, and HP_010, is a known virulence factor that counteracts stomatal closure, aiding entry into the host (Schellenberg et al., 2010). Cichofactin, present in HP_013, is a lipopeptide that has demonstrated roles in virulence, swarming motility, and biofilm formation in *P. cichorii* (Pauwelyn et al., 2013). Finally, viscosin, found only in HP_015, is a lipopeptide biosurfactant that has been shown to enhance biofilm dispersal in *P. fluorescens* (Bonnichsen et al., 2015). Together, the results of virulence-associated genomic characterisation suggest many of these isolates have pathogenic potential, especially if they are acting collaboratively with pathogens like Psa which carries a large effector repertoire itself (McCann et al., 2013, 2017). The presence of PAIs, effectors, phytotoxins and other secondary metabolites suggest that some of the isolates could have contributed to disease observed in the kiwifruit germplasm. The diverse secondary metabolite profiles of these isolates are of particular interest: they may aid Psa in its infection process, as local Psa3 isolates possess only a putative NRPS biosynthetic toxic cluster, unlike other Psa biovars which carry toxins such as coronatine and phaseolotoxin (Fujikawa & Sawada, 2019; Sawada & Fujikawa, 2019).

Conversely, we also observed isolates that reduced the growth of Psa3 V-13 when co-inoculated, namely *P. syringae* phylogroup 3 isolate HP_007 and *P. protegens* isolate HP_009. *P. protegens* isolates have been shown to inhibit bacterial phytopathogens (Gu et al., 2022; Ortega et al., 2020), with HP_009 encoding secondary metabolite pathways associated with plant-protecting *P. protegens* strains, including 2,4-diacetylphloroglucinol (DAPG), pyoluteorin, and hydrogen cyanide (Gu et al., 2022). DAPG is a broad-spectrum antibiotic produced by *Pseudomonas* spp. with disease-suppressing activity (Johnson et al., 2023). Secondary metabolites with antimicrobial activity were also observed in other isolates, such as bananamide which has selective anti-fungal activity (Omoboye et al., 2019); rhizomide A, a rhizomide-class lipopeptide, which has weak anti-oomycete and antibacterial activity (X. Wang et al., 2018); and nunapeptin/nunamycin, which has antimicrobial activity (Hennessy et al., 2017).

Co-inoculation results suggest that some *Pseudomonas* species may be able to aid each other to increase proliferation in multiple species of *Actinidia.* Psa alone was unable to cause symptoms on resistant germplasm hosts. Similarly, individual germplasm isolates were also non-pathogenic on *A. arguta, A. polygama* and *A. melanandra* hosts when infected alone, as indicated by their inability to accumate to a ‘disease’ level such as that seen for Psa in susceptible ‘Hort16A’. Host-dependent growth trends were still observed, unsurprisingly given that host compatibility is a central part of pathogen success (Sacristan & Garcia-Arenal, 2008). The use of Pfm as a control strain exemplifies the host-specific nature of interaction outcomes. When co-inoculated with Psa3 V-13, Pfm exhibited mutualistic pathogenicity in *A. arguta,* suppression of Psa3 V-13 growth in *A. melanandra,* and no change in growth on *A. polygama*. Notably, Pfm has all the makings of a successful pathogen, including a T3SS and large effector repertoire, yet it is unable to infect both susceptible and resistant hosts alike (Jayaraman et al., 2023). Many factors may affect this compatibility, including effector and toxin repertoires, T3SS presence and type, and host immune responses. The disease symptoms observed in these co-inoculation assays across biological replicates also suggests the possibility of pathogen complex formation in the *Actinidia* phyllosphere. This phenomenon is known to occur in other hosts. For example, synergism has been demonstrated between *P. cichorii, P. corrugata, P. viridiflava, P. mediterranea, P. fluorescens, Pectobacterium* spp. and *Dickeya chrysanthemi* to cause tomato pith necrosis (Lamichhane & Venturi, 2015). *Xanthomonas perforans* was shown to exhibit higher disease severity in co-infection with *X. arboricola* and *P. capsici* than alone, or dually with either species (Sadhukhan et al., 2024). Consortia formation has also been observed between *P. savastanoi* pv. *savastanoi* (Psv), *Pantoea agglomerans,* and *Erwinia toletana* (Hosni et al., 2011). *P. agglomerans* is particularly interesting in that it can suppress or aid Psv knot formation (Hosni et al., 2011). This suggests that commensal bacteria are flexible in their interactions with pathogens, with pathogenic consortia neatly exemplifying the breadth of the symbiosis spectrum. These interactions are often facilitated by the sharing of virulence factors and secondary metabolites (Sadhukhan et al., 2024). Co-operative virulence through public goods sharing was also recently demonstrated using a *P. syringae* pv. *tomato* (Pto) metaclone, where Pto effectors were individually cloned onto plasmids, transformed into an effectorless background and mixed to create a ‘metaclone’ (Ruiz-Bedoya et al., 2023). When plants were co-inoculated with the virulent metaclone and non-virulent *Pseudomonas fluorescens,* the growth of *P. fluorescens* increased substantially (Ruiz-Bedoya et al., 2023). This further suggests that pathogenic species may confer the benefits of virulence to unrelated members of the phyllosphere, and that even non-virulent members of the phyllosphere could benefit opportunistically. Although pathogen effectors can act as public goods, this work also suggests that host specificity plays an important role in determining co-operative outcomes. These results add to a growing body of research that shows bacteria can cause disease socially, often with greater severity (Sadhukhan et al., 2024). Pathogenic consortia may have consequences for the epidemiology of Psa incursions, and may further complicate the diagnosis and subsequent control of complex multispecies diseases (Lamichhane & Venturi, 2015).

Conversely, other isolates reduced the growth of Psa *in planta*. Suppressive microbial communities have been demonstrated in a number of systems. For example, closely related commensal *Pseudomonas* spp. have been demonstrated to suppress pathogenic *Pseudomonas spp.* (Wang et al., 2022). Similarly, phyllosphere bacterial consortia from field-grown soybeans can reduce the disease symptoms of *P. syringae* pv. *glycinea* in soybean, benefiting plant growth and health (Agbavor et al., 2022). Ehau-Taumaunu & Hockett (2022) have demonstrated that bacteriocin-mediated antagonism increases the Pss population up to eight-fold *in planta* when co-infiltrated with the bacteriocin-sensitive *P. syringae* pv. *phaseolicola*. Furthermore, this bacteriocin-conferred growth benefit was not observed *in vitro* and appears to be dependent on the virulence conferred by effector secretion (Ehau-Taumaunu & Hockett, 2022). This again highlights that pathogen proliferation relies on suppression of host defence responses and forms the background against which microbe-microbe interactions take place. Disease-suppressive phyllosphere communities have also been successfully developed through serial passaging, reducing disease severity in tomato infected with Pto DC3000 (Ehau-Taumaunu & Hockett, 2023). It is worth noting that the ratio at which these isolates were co-inoculated is not reflective of the abundance of these isolates in the field. Straub et al. (2018) performed competitive growth assays with Psa3 NZ54 and the *P. syringae* phylogroup 3a isolate G33C at both a 1:1 ratio and a ‘rare’ invasion ratio of 1:100 on *A. chinensis* var. *chinensis* varieties. Co-inoculation reduced the epiphytic and endophytic growth of Psa in early growth stages, in a variety-specific manner (Straub et al., 2018). On the other hand, co-inoculation increased the growth of *P. syringae* G33C, even when co-inoculated with a T3SS-lacking Psa3 Δ*hrcC* mutant which cannot secrete effectors, suggesting that this benefit cannot be attributed solely to Psa’s effector-mediated virulence (Straub et al., 2018). The effects of population density on colonisation, resource competition, antimicrobial compound and phytotoxin accumulation, contact-dependent growth inhibition, and bacteriocin action must be considered (Straub et al., 2018). Future work should endeavour to build synthetic communities that better represent in-field communities.

The combined or collective pathogenicity of isolates in the *Actinidia* phyllosphere does not necessarily depend on virulence-associated features, such as effectors and phytotoxins, of the co-infecting partner. There did not appear to be any strong correlations between virulence-associated factors and observed pathogenicity in this work. On the contrary, we have demonstrated that commensal or weakly pathogenic bacteria can move across the spectrum of symbiosis and contribute to disease, similarly to interactions between *Pseudomonas* spp. observed in *Phaseolus vulgaris* (Rufián et al., 2018). Isolates with small effector and toxin repertoires significantly increased *in planta* growth during co-inoculation, whilst isolates like HP_006 with relatively larger repertoires had no effect on Psa growth *in planta*. Strains in *P. syringae* phylogroup 2 are thought to possess fewer effectors than other phylogroups, while maintaining conserved toxin pathways, possibly making them better epiphytes than other phylogroups (Hockett et al., 2014). It is possible that these smaller repertoires allow these isolates to manage the burden of Psa’s recognition, allowing proliferation of both the isolate and Psa3 V-13. The results of this work, particularly the possible formation of pathogen complexes, are concerning. However, this work demonstrates a range of interaction outcomes, with other isolates showing the potential to reduce Psa’s growth. Further testing is needed to determine how isolates might interact in more complex community assemblages *in planta.* Future work should also explore extending this view of microbial interplay, to capture fungi and other eukaryotic microbes present in the phyllosphere.

Finally, recent advances in metagenomics could help us understand the prevalence of species, lineages, and variants in the disease-associated phyllosphere (Marbouty et al., 2014; Burton et al., 2014; Bickhart et al., 2022; Benoit et al., 2024). Metagenome-assembled genomes (MAGs) may allow for less-biased detection of potential and emerging pathogens, given the labour-intensive nature of traditional isolation-based genome biosurveillance. Further still, metagenomics offers a powerful lens to improve our understanding of phyllosphere communities, allowing us to identify and track effector repertoires across all members, providing insights into the potential for the emergence of virulence or co-operative phenotypes. Alternatively, this could also be achieved through targeted amplicon sequencing of genes of interest, particularly effectors and known toxins. This approach has recently been demonstrated Byers et al. (2024), who surveyed the of diversity of secondary metabolite biosynthetic gene clusters in kauri forest soils. By characterising the genomic potential of these communities, we can better understand the broader context in which known pathogens like Psa exist, and from which both new pathogens and new biocontrol agents may emerge. The research presented here provides a foundation for future studies aimed at elucidating the mechanisms of co-operative virulence and disease suppression in the phyllosphere, helping to protect crops against emerging threats, potentially through the development of new disease management strategies and agents.

## Methods

### Germplasm sampling

Two kiwifruit germplasm collections held at Plant & Food Research’s Kerikeri and Te Puke Research Orchards were sampled. Each comprised multiple blocks of diverse, unimproved *Actinidia* spp. vines, grown from kiwifruit seeds collected from the wild in China (Richardson et al., 2023). Symptomatic leaves were collected from Te Puke and Kerikeri in December 2022. One hundred leaves were taken from Te Puke, and 133 leaves were taken from Kerikeri. At least two leaves were sampled per vine, and vines were selected for sampling to represent all *Actinidia* species displaying leaf spot symptoms. Hands and tools were sanitised using 80% ethanol between vines. Leaves sampled at Te Puke were stored in insulated boxes with ice packs for transport back to Plant & Food Research’s Mount Albert Research Centre (MARC) and held overnight at 4°C before processing the next day. Leaves sampled at Kerikeri were kept ambient during sampling and stored at 4°C overnight at Kerikeri. They were then transported back to MARC in insulated cold boxes and stored overnight at 4°C before processing the next day. Samples were processed 1-2 days post-harvest. Leaves were photographed and the features of visible lesions were noted. Leaf homogenate was prepared from all leaves sampled at Te Puke, totalling 100 symptomatic samples and 32 asymptomatic samples. Only a selection of leaves sampled from Kerikeri were processed to avoid duplicate genotype representation. The Kerikeri leaf samples included 99 symptomatic and 87 asymptomatic samples.

Symptomatic leaf discs were taken from visible lesions, while asymptomatic leaf discs were sampled from visually asymptomatic portions of the same leaf. For each sample, up to four leaf discs were taken using a 5 mm cork-borer either from symptomatic or asymptomatic portions of leaf. Leaf discs were immediately submerged in 350 μL sterile 10 mM MgSO_4_ with three stainless steel beads. Tools and surfaces were sterilised with 70% ethanol between each sample. Samples were homogenised using a Storm24 Bullet Blender (Next Advance, New York, USA) at speed twelve for 2 min. An additional 350 μL of MgSO_4_ was added and the homogenisation was repeated. Leaf homogenate was stored at −20°C prior to DNA extraction.

### Amplicon sequencing

#### Sample selection

A total of 74 samples were selected for amplicon sequencing: 37 symptomatic and 37 corresponding asymptomatic samples (Supplementary Table 4). Samples were selected to represent every *Actinidia* species sampled, with equal representation across both sites where possible. Priority was given to species that are retained in tissue culture by Plant & Food Research, for downstream continuity of *in planta* assays. These included *A. polygama, A. melanandra,* and *A. arguta*. Within these species, a range of genotypes were selected, where possible. Two sterile tissue culture samples (*A. arguta* AA07_03 and *A. polygama* PC02_01) were also included in the amplicon sequencing as negative controls along with a sterile water non-template control. Along with the phyllosphere samples, we included a ZymoBIOMICS Microbial Community Standard (Zymo Research, California, USA) as a ‘mock’ community control for read depth and accuracy, and a sample of *P. syringae* pv. *actinidiae* V13 genomic DNA as a positive control.

#### DNA extraction

A QIAGEN DNeasy® Plant Mini Kit (QIAGEN) was used to extract DNA from leaf samples for 16S amplicon sequencing. The kit was used according to the manufacturer’s provided quick-start protocol. Non-template controls were used for all extractions and later were screened for contamination with qPCR utilising Psa-specific and universal bacterial ITS primers. DNA was then stored at 4°C.

#### Quantitative PCR

Quantitative PCR was used to ascertain bacterial presence in the samples selected for amplicon sequencing, and to test for the presence of Psa. Psa F1/R2 primers specific to Psa and Pfm were used, (Rees-George et al., 2010), as well as ‘universal’ 335F/769R primers for amplifying bacterial 16S rRNA gene regions (Supplementary Table 5; Dorn-In et al., 2015). Plant reference gene (EF1α) primers, SN126L/R (Nardozza et al., 2013), were used for normalising results. The primers used to identify nonspecific bacterial presence by qPCR, known as 335F/769R, were shown to have an efficiency of 102% and an amplification factor of 2.02. Non-template controls were used for all runs with each primer pair, with wild-type (WT) Psa3 V-13 genomic DNA (gDNA) used as a positive control. Quantitative PCR (qPCR) was performed using an Illumina Eco™ Real-Time PCR System. Quantitative PCR parameters were based on previous work by Andersen et al. (2018).

#### End-point PCR and sequencing

End-point PCR was performed in 50 μL reactions in 0.2 mL PCR tubes. Master-mixes were made fresh for each run and aliquoted into tubes, followed by template DNA (Supplementary Table 6). End-point PCRs were conducted in an Eppendorf Mastercycler X50I thermal cycler (Eppendorf, Hamburg, Germany) using the parameters outlined in Supplementary Table 7. Symptomatic phyllosphere samples were amplified for 30 PCR cycles, and asymptomatic samples were amplified for 40 PCR cycles. The ZymoBIOLOMICS Microbial Community Standard and Psa3 V-13 gDNA controls were amplified for 30 cycles. The no-template control (NTC), a ‘blank’ sample that went through all extraction processes, was amplified for 40 cycles.

Samples were amplified (using the conditions listed above) with the 16S rRNA V5-V9 primers 799F (AACMGGATTAGATACCCKG) and 1492R (TACGGYTACCTTGTTACGACTT), generating a ∼690 bp product (Heuer et al., 1997; Chelius & Triplett, 2001). Samples were prepared with AMPure XP (Beckman Coulter Life Sciences, Washington, USA) bead clean-up. Amplicon concentration was quantified with Qubit (Thermo-Fisher Scientific, California, USA) and normalised to ∼5 ng/μL.

Nanopore sequencing was performed using an SQK-LSK114 Ligation Sequencing Kit with EXP-PBC096 barcoding on a PromethION FLO-PRO114M flowcell through Auckland Genomics (Auckland, New Zealand). Real-time base-calling was done using a high-accuracy model with Guppy 7.1.4. Adaptors were trimmed but barcodes were not. After the run was complete, the reads were “re-base-called” using a Super High accuracy model with Dorado 0.5.3 and model dna_r10.4.1_e8.2_400bps_sup@v4.2.0. Quality checking of sequences was performed with both FastQC (Andrews, 2010) and MultiQC (Ewels et al., 2016).

#### Bioinformatic & statistical analysis

Data analysis was conducted in R (version 4.2.1; R Core Team, 2024). 16S amplicon sequencing reads were processed into amplicon sequence variants (ASVs) using the R package dada2 (version 1.24.0; Callahan et al., 2016). ONT reads were treated as single-end reads and were filtered for quality, using relaxed maxEE values (maxEE=15, truncQ=2, minLen=400, maxLen=1000). Taxonomy was assigned to the genus level using the SILVA v132 database for both forward and reverse-complement orientations (tryRC=TRUE, minBoot=80), with further species-level assignments made based on exact matching to reference sequences. The R package decontam was used to identify contaminant ASVs for removal which were present in no-template control samples (version 1.18.0; Davis et al., 2018). The R package phyloseq (version 1.42.0; McMurdie & Holmes, 2013) was used to create a phyloseq object of metadata and count data, perform ordination, and visualise results. Each sample was normalised to the median sequencing depth (https://joey711.github.io/phyloseq/preprocess.html). Taxonomy was visualised at the genus and species levels (where applicable) using the plot_bar() function (McMurdie & Holmes, 2013). The merge_samples2() function from the speedyseq package was used to visualise relative abundance by data levels (version 0.2.0; https://github.com/mikemc/speedyseq). Alpha and beta diversity were calculated as laid out in the meta-analysis pipeline from Hoffbeck et al. (2023). The plot_richness() function was used to calculate alpha-diversity using the Shannon diversity metric (McMurdie & Holmes, 2013). The ordinate() function was used to calculate beta-diversity using a Bray-Curtis dissimilarity matrix (McMurdie & Holmes, 2013). PERMANOVA statistical significance were calculated using Bray-Curtis dissimilarity with 9999 permutations using the adonis() function from the vegan package (version 2.6-2; Oksanen et al., 2022). *Pseudomonas* ASVs were identified to the species or sub-species level through BLASTn and comparison to reference sequences (Johnson et al., 2008). The core_members() function from the microbiome package was used to identify core members of the microbiome (80% core threshold; version 1.20.0; Lahti & Shetty, 2017).

### Whole genome sequencing

#### Sample selection

Symptomatic leaf homogenate (200 μL) was streaked onto 60 mm Lysogeny Broth (LB) agar (LBA) plates containing 25 μg/mL nitrofurantoin and 40 μg/mL cephalexin and incubated at room temperature. When colonies could be observed, each plate was noted for fungal presence and bacterial morphology. *Pseudomonas* -like colonies were then selected from each plate and passaged three times to obtain pure culture. If *Pseudomonas* -like morphology was not seen, the predominant colony type was selected. After the third passage, single colonies were inoculated into 2 mL of LB broth and incubated at room temperature for two days with shaking at 200 rpm. Liquid culture (500 μL) was then added to an equal volume of 50% glycerol and frozen at −80°C. One mL of the liquid inoculum was also retained for DNA extraction. Seventy μL of the crude leaf homogenate from both the symptomatic and asymptomatic samples was inoculated into 2 mL of non-selective LB broth and incubated for two days at room temperature with shaking at 200 rpm. After incubation, 500 μL of the liquid culture stored at −80°C as described previously.

Samples were selected for whole genome sequencing after the third passage stage described above. As several non-Psa colony phenotypes were observed, isolates were selected to represent a variety of these phenotypes as well as sampling for representation across regions and host genotypes (Supplementary Table 8).

#### DNA extraction and sequencing

A Promega Wizard® Kit (Promega Corporation, Wisconsin, USA) was used to extract DNA for whole genome sequencing. The kit was used according to the manufacturer’s instructions, with the exception that 3 μL of RNAse was rather than the recommended 2 μL.

Whole genome sequencing was performed by Novogene China (https://www.novogene.com) via Auckland Genomics (https://www.auckland.ac.nz/en/science/about-the-faculty/share-shared-research-equipment/auckland-genomics.html). Library preparation was done using purePlex™ DNA Library Prep Kit (seqWell, Beverly, Massachusetts, USA). Sequencing was carried out using an Illumina MiSeq 2x150bp PE Nano platform.

#### Bioinformatic analysis

Whole genome sequencing reads were assembled using shovill (version 1.1.0; https://github.com/tseemann/shovill) and the resulting contigs were annotated with Prokka (version 1.14.5; Seemann, 2014) using the default prokaryotic database. Reference isolates were also annotated with Prokka (Seemann, 2014). The phylogeny was built on representative isolates from the *Pseudomonas syringae* species complex phylogroups from Dillon et al., (2019), as well as non-syringae representatives from each *Pseudomonas* subgroup. These isolates were used for pangenome analysis with panaroo (version 1.3.0; Tonkin-Hill et al., 2020) to produce an alignment of core genes (panaroo options: --clean-mode strict -a core --remove-invalid-genes --aligner mafft --core_threshold 0.90). A phylogeny was produced from this core gene alignment with RAxML (options -f a -p 12345 -x 12345 -# 100 -m GTRCAT version 8.2.10; Stamatakis, 2014). The R package ggtree (Yu et al., 2017) was used to visualise the resulting phylogenies.

Magic-BLAST (version 1.7.2; Boratyn et al., 2019) was used to BLAST custom databases of reference type III secretion systems (T3SSs) and effectors against germplasm isolates. The databases for both effectors and T3SSs were created with sequence data from Dillon et al. (2019) using the magic-BLAST function ‘makeblastdb’ (options: -dbtype nucl -parse_seqids; Boratyn et al., 2019). The databases were then used to search against the germplasm isolates using the NCBI+ function blastn (Camacho et al., 2009). To validate effector hits, putative effectors were mapped to germplasm isolate genomes manually in Geneious Prime (version 2022.0.1), and the best alignment was selected. For T3SS analysis, homologous regions were extracted from representative genomes to build a custom database (Supplementary Table 9). For *P. viridiflava* RMX3.1b and *P. viridiflava* PNA3.3a, the PAI region was downloaded directly from NCBI as identified and annotated by Araki et al. (2006). The database was then used to search the germplasm isolates to identify the most similar T3SS. Once a candidate T3SS was identified, the individual genes of the corresponding T3SS were mapped to the germplasm isolate genome. Toxins were identified using antiSMASH under relaxed detection strictness (version 7.0; Blin et al., 2023). Results with under 60% similarity were then filtered out.

### In planta assays

Germplasm-derived isolates were selected from the isolates which underwent whole genome sequencing. They were selected to represent a range of virulence associated genomic features present, sampled regions, and taxonomic and phylogenetic diversity if within the *P. syringae* species complex (Table 1). Psa3 V-13 (ICMP 18884) and Pfm LV-5 (ICMP 18803) were used as controls.

**Table 1.** Table with details for germplasm-derived isolates used in pathogenicity testing with *Pseudomonas* syringae pv. *actinidiae* biovar 3 (Psa3) V-13.

| Isolate | Closest relative | Proposed phylogroup | Host | Region |
| --- | --- | --- | --- | --- |
| HP_001 | *P. graminis* | - | *A. arguta* | Te Puke |
| HP_004 | *P. cichorii* | 11 | *A. arguta* | Te Puke |
| HP_006 | *P. syringae* | 2 | *A. melanandra* | Te Puke |
| HP_007 | *P. syringae* | 3 | *A. melanandra* | Te Puke |
| HP_009 | *P. protegens* | - | *A. arguta* | Te Puke |
| HP_013 | *P. viridiflava* | 7 | *A. arguta* | Kerikeri |
| HP_014 | *P. quasicaspiana* | 13 | *A. polygama* | Kerikeri |
| HP_015 | *P. sivasensis* | - | *A. polygama* | Kerikeri |

Tissue culture plantlets were used for the inoculation of bacterial isolates. All plantlets originated from Plant and Food Research orchard-sourced spring budwood that had been transferred into tissue culture. *Actinidia arguta* AA07_03 plantlets were sourced from Multiflora (Auckland, New Zealand) who maintain and propagate the tissue culture. *A. melanandra* and *A. polygama* plantlets were propagated in-house at Plant & Food Research’s MARC (Auckland, New Zealand). All plantlets were grown in 400 mL lidded plastic pottles on half-strength Murashige and Skoog (MS) agar. All pottles contained between 3 and 5 cuttings and were only used once sufficient leaf surface area for harvesting twenty 1-cm leaf discs was observed.

#### Pathogenicity assays

The pathogenicity of bacterial strains was assessed via *in planta* flooding inoculation as per McAtee et al., (2018). Bacterial strains were grown from frozen glycerol stocks on LB plates supplemented with nitrofurantoin and cephalexin. Colonies were then selected and incubated overnight in 5 mL liquid LB with shaking at 250 rpm at approximately 20°C. After incubation, 2 mL of culture was centrifuged at 5,000*g* for 3 min and the supernatant replaced with 1 mL 10mM MgSO_4_. The resuspended bacterial cultures then acclimated for 1 h at room temperature, before reading the optical density of cells at 600nm (OD_600_). Inocula were then diluted in 500 mL 10mM MgSO_4_ to approximately 10^6^ CFU/mL (OD_600_=0.005) and Silwet™ L-77 (0.0025%; Momentive Performance Materials, New York, USA) was added. Inoculation was carried out by submerging kiwifruit plantlet containers with the bacterial cells in 500 mL of 10mM MgSO_4_ for approximately 3 min. After 3 min, the inoculum was decanted, and the plantlet containers were resealed for incubation at 20°C with a 16h:8h light:dark cycle. Leftover inoculum (50 μL) was plated at 1000x dilution to validate viability. Competition assays were conducted the same way with the exception that cultures were combined in a 1:1 ratio prior to flooding. Co-inoculated isolates were combined at the same final concentrations.

Leaf samples were taken at 7 days post-inoculation. Four technical replicates were taken per plantlet, with four 1-cm leaf discs per technical replicate. Competition assays (germplasm isolate + Psa3 V-13) were performed with three biological replicates. Leaf discs were washed in sterile MilliQ water for 30 s, and each technical replicate of four leaf discs placed into an Eppendorf™ Microcentrifuge Safe-Lock™ tube (Eppendorf, Hamburg, Germany) with three sterile 3.5mm stainless steel beads and 350 μL sterile 10 mM MgSO_4_. Samples were macerated for two rounds of one min each with manual dislodgement of any pellets formed between rounds of maceration at top speed in a Storm24 Bullet Blender (Next Advance, New York, USA).

Leaf homogenate was then serially diluted to 10^-5^ and the diluted homogenate plated out on LBA plates containing 25 μg/mL nitrofurantoin and 40 μg/mL cephalexin. Colonies were then counted at the lowest dilutions possible and used to calculate CFUs/cm^2^. DNA was extracted from the leaf homogenate using a QIAGEN DNEasy Plant Mini Kit (QIAGEN, Hilden, Germany), according to the manufacturer’s instructions. Extracted DNA was then assessed for bacterial growth using qPCR.

#### Quantitative PCR

Quantitative real-time PCR was used to quantify the results of pathogenicity assays in addition to or in place of plate count data. The primers used for each germplasm isolate are outlined in Supplementary Table 10. The primers were validated against the target isolate using PCR and subsequent gel visualisation, as well as against Psa3 V-13 as a negative control. The primers were then tested using a serial dilution of the target isolates to assess primer efficiency and all were assessed to be within the valid range for use (90-120% efficiency). qPCR parameters were based on previous work by Andersen et al. (2018).

#### Data visualisation

Analysis of pathogenicity assay data was performed in R (version 4.2.1; R Core Team, 2024). The packages ‘tidyverse’ (version 2.0.0; Wickham et al., 2019) and ‘ggplot2’ (version 3.5.1; Wickham, 2016) were used for data manipulation and plot creation, respectively. Normality of data was assessed using Shapiro-Wilks test. For comparisons between individual- and co-inoculations, the function stat_compare_means() from the ggpubr package (version 0.6.0; Kassambara, 2023) was used. Kruskal-Wallis and Wilcoxon tests were used to analyse statistical differences between groups for non-normal data; one-way ANOVA and Welch’s t-test were used to compare normally distributed data. Treatments were pairwise compared with Tukey’s honestly significant difference (HSD) post hoc at 5% and compact letter display was used on plots.

## Acknowledgements

The authors thank Sidney Scott for assistance with germplasm sampling, Juanita Dunn and Gavin Lloyd for assistance with research orchard access, and Drs Hanareia Ehau-Taumaunu, Virginia Marroni and Hayley Ridgway for manuscript review. We thank Dr Nikki Freed & the team at Auckland Genomics for DNA sequencing support. The authors also acknowledge the use of New Zealand eScience Infrastructure (NeSI) high-performance computing facilities, consulting support and training services as part of this research. New Zealand’s national facilities are provided by NeSI and funded jointly by NeSI’s collaborator institutions and through the Ministry of Business, Innovation & Employment’s Research Infrastructure programme. URL https://www.nesi.org.nz.

## Data availability statement

Sequence data are available in the GenBank Nucleotide Database (https://www.ncbi.nlm.nih.gov/genbank/) and Sequence Read Archive (https://www.ncbi.nlm.nih.gov/sra) under BioProject PRJNA1213159.

## Author contributions

Haileigh R. Patterson: Data curation; Formal analysis; Investigation; Methodology; Project administration; Software; Validation; Visualisation; Writing – original draft; Writing – review & editing.

Lauren M. Hemara: Conceptualisation; Data curation; Formal analysis; Investigation; Methodology; Project administration; Software; Supervision;

Validation; Visualisation; Writing – original draft; Writing – review & editing.

Jay Jayaraman: Conceptualisation; Funding acquisition; Methodology; Project administration; Supervision; Writing – review & editing.

Matthew D. Templeton: Data curation; Funding acquisition; Supervision; Writing – review & editing.

## Supporting information

**Supplementary Figure 1.**
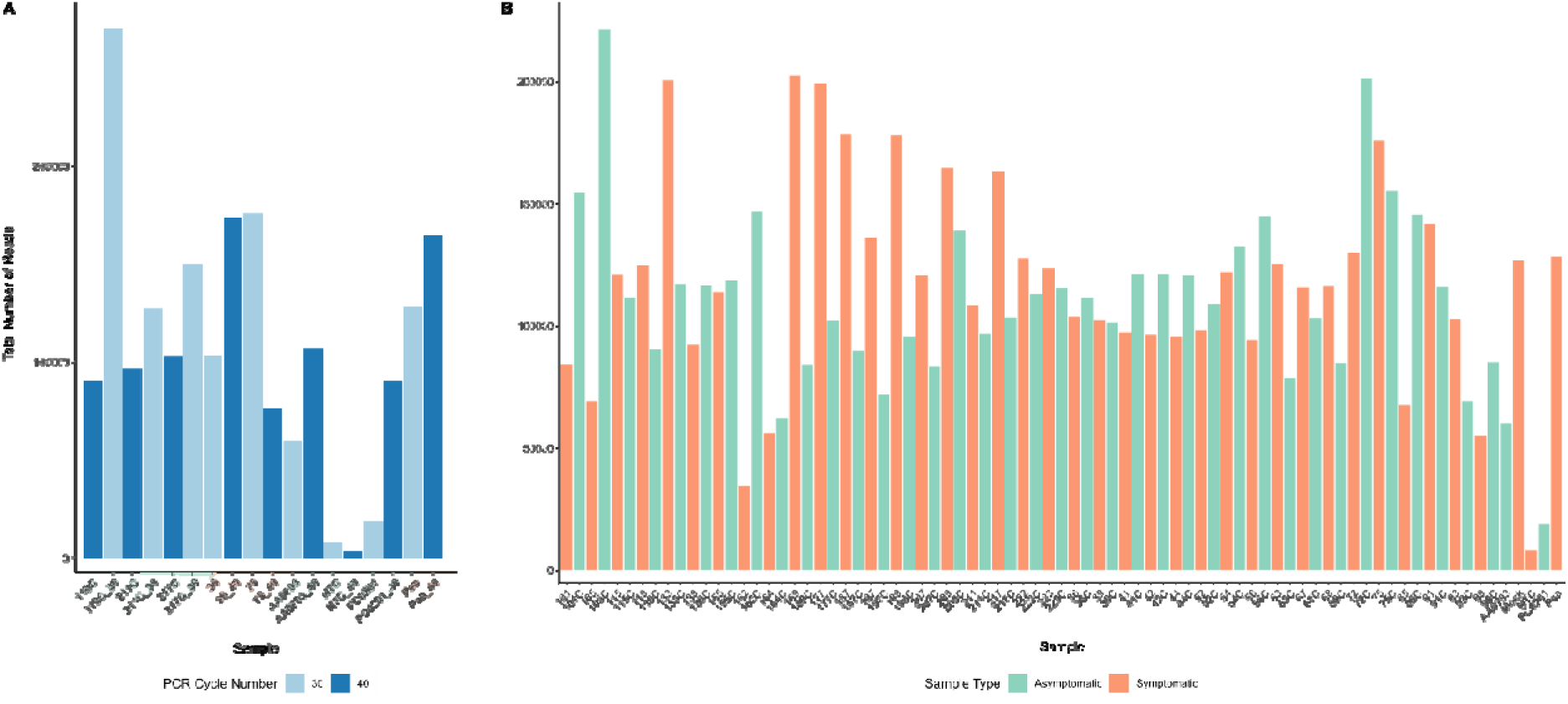
Total number of 16S amplicon sequencing reads in paired controls and all samples. Total number of sequencing reads was compared for (A) samples that were amplified at both 30 and 40 PCR cycles with sample type indicated with colour highlighting on the x-axis, and (B) symptomatic and asymptomatic sample pairs. The no-template control (NTC) was a ‘blank’ sample that went through all extraction and amplification processes. AA0703 and PC0201 samples were taken from axenic tissue culture plantlets. The ‘Psa’ samples are Psa3 V-13 gDNA.

**Supplementary Figure 2.**
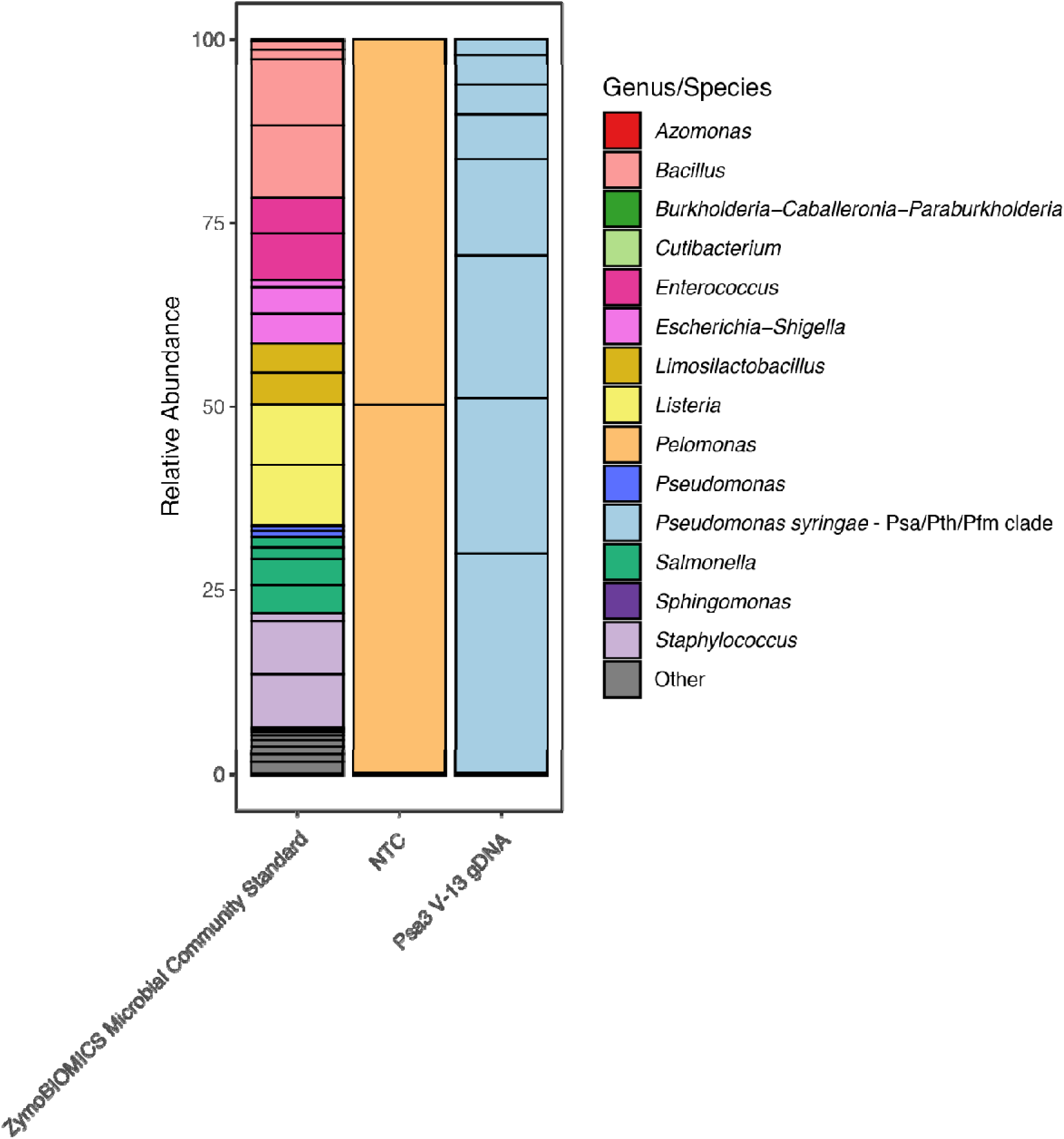
Dominant bacterial genera present in amplicon sequencing control samples. The 50 most abundant amplicon sequence variants (ASVs) are represented, coloured by genus. ASVs identified as ‘*Pseudomonas syringae* – Psa/Pth/Pfm clade’ were identified by BLASTn. All other taxa are indicated by the group ‘Other’. Each black box within the bar indicates a different ASV. The ZymoBIOLOMICS Microbial Community Standard and *Pseudomonas syringae* pv. *actinidiae* biovar 3 (Psa3) V-13 gDNA controls were amplified for 30 cycles. The no-template control (NTC) was a ‘blank’ sample that went through all extraction and amplification processes, which was amplified for 40 cycles.

**Supplementary Figure 3.**
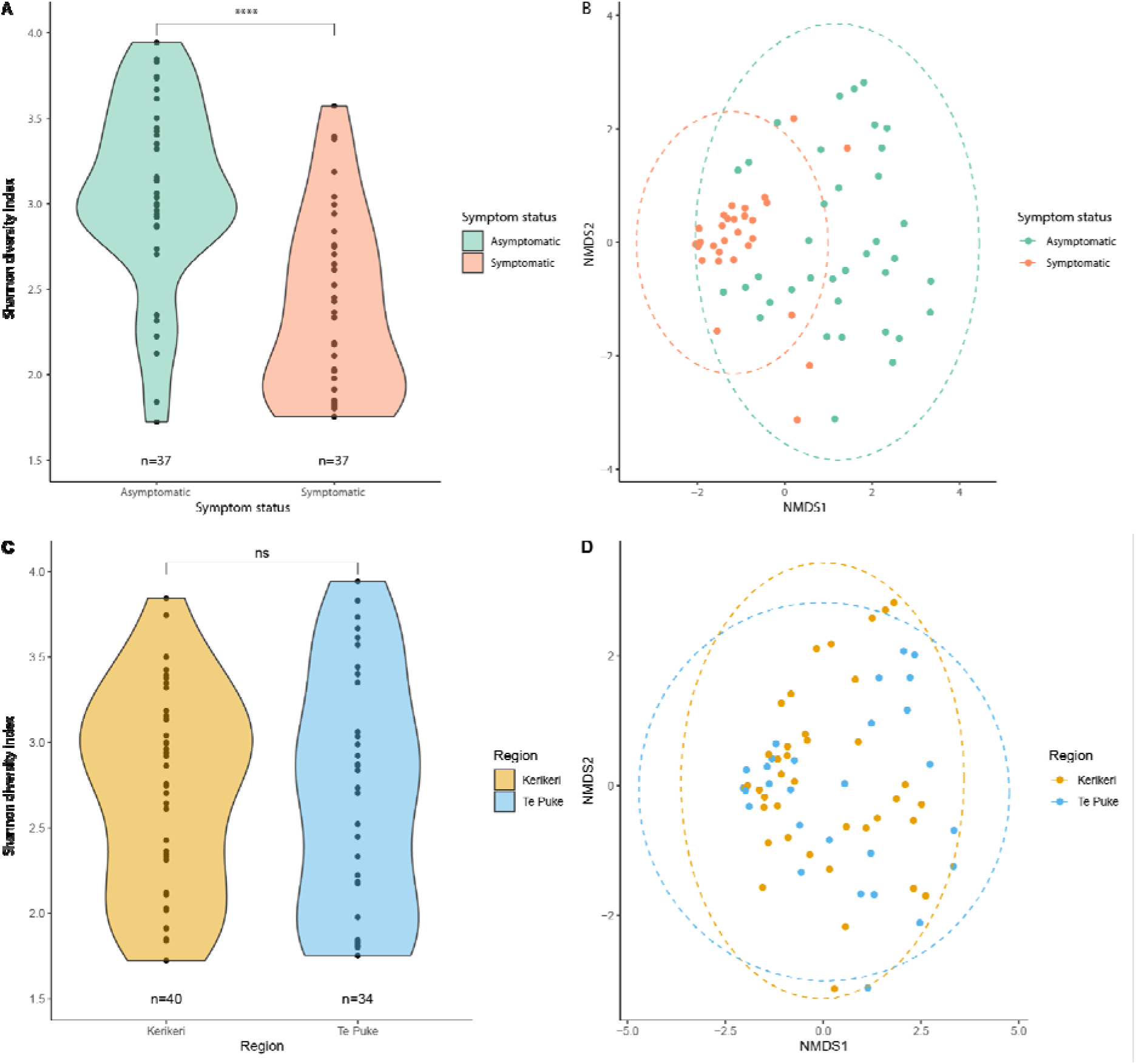
Bacterial community alpha-diversity and composition by symptomatic status and region. Bacterial alpha-diversity using the Shannon metric by (A) symptom status and (C) region. Points represent individual samples. Asterisks indicate the statistically significant difference of Wilcoxon’s test, where *p*≤0.0001 (****) and *p*≥0.05 (ns). Non-metric multidimensional scaling (nMDS) ordination showing the effect of (B) symptom status and (D) region on bacterial community composition. Ellipses represent statistically similar communities with a 95% confidence interval.

**Supplementary Figure 4.**
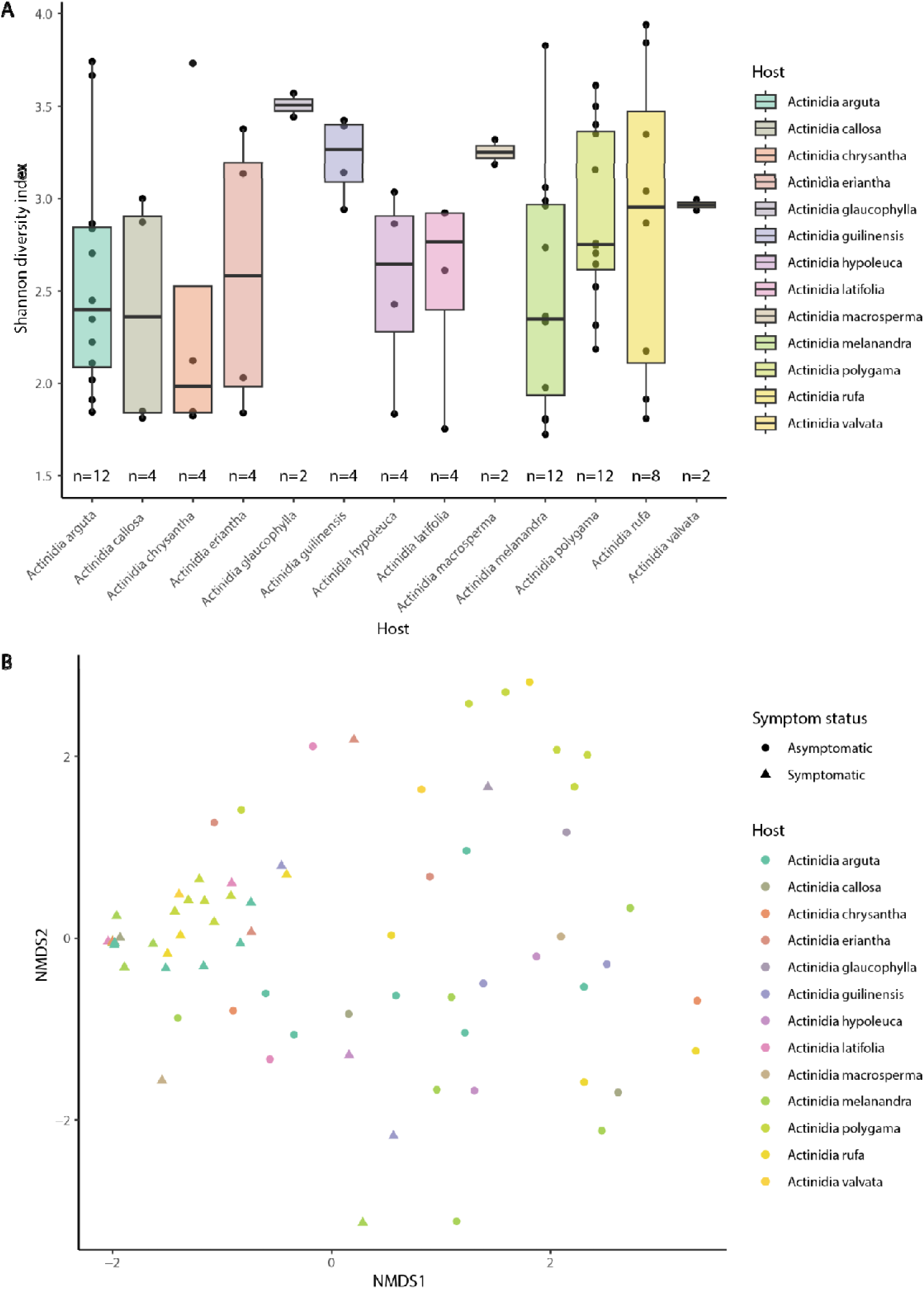
Bacterial community alpha-diversity and composition by host species. (A) Bacterial alpha-diversity using the Shannon metric by host. Points represent individual samples.(B) non-metric multidimensional scaling (nMDS) ordination showing the effect of host on bacterial community composition.

**Supplementary Figure 5.**
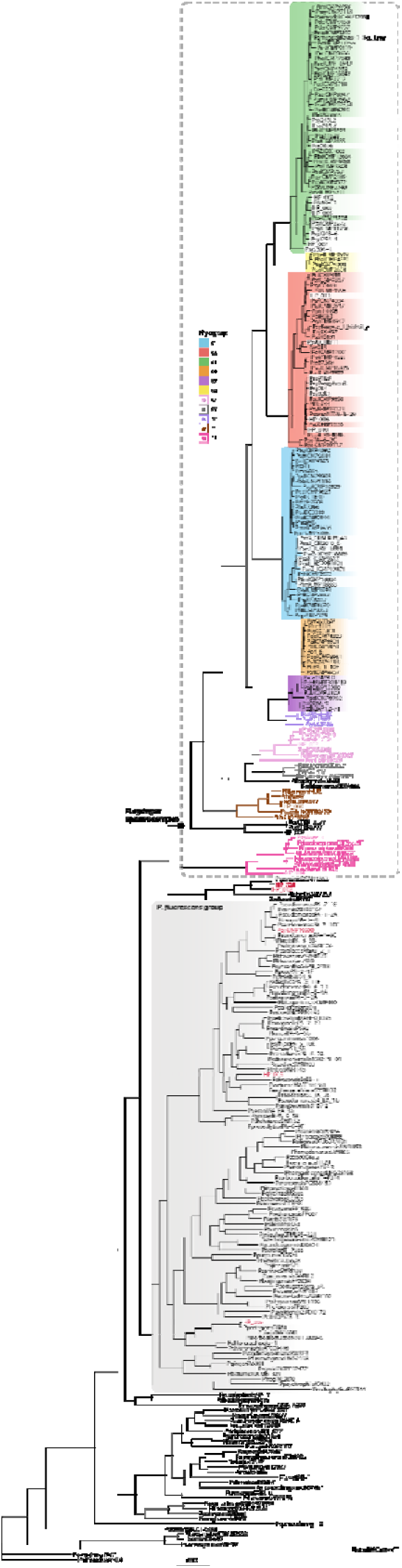
Core gene phylogeny of germplasm isolates, representative *Pseudomonas syringae* isolates, and plant-associated *Pseudomonas* species. A core gene phylogeny was produced with Panaroo (version 1.3.0; Tonkin-Hill et al., 2020) and RAxML (version 8.2.12; Stamatakis 2014), with a core gene threshold of 90%. The *P. syringae* species complex (indicated by the grey dashed box) is coloured by phylogroup, as designated by Dillon et al. (2019). Isolates from *Actinidia* spp. are bolded and highlighted.

**Supplementary Figure 6.**
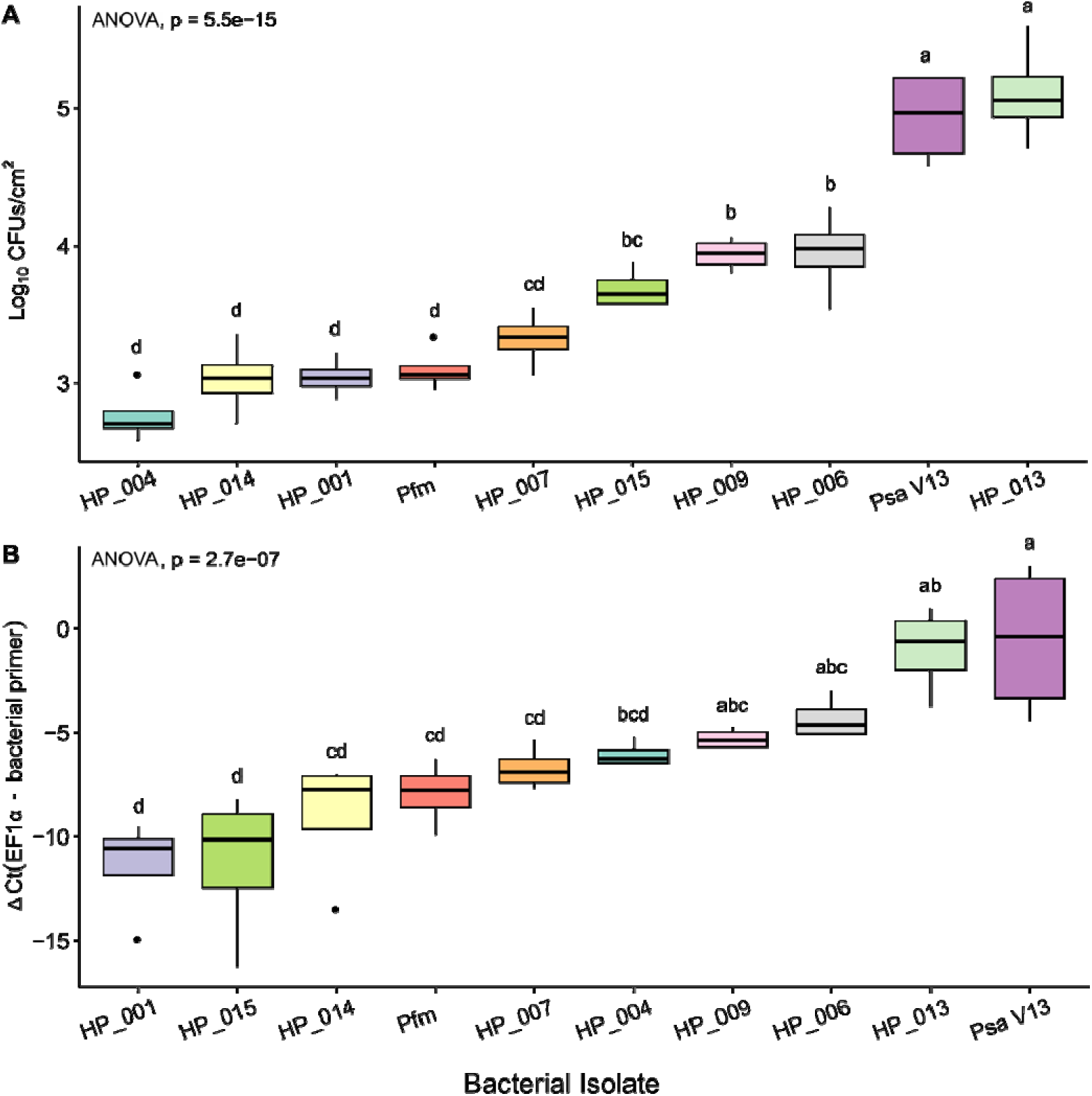
Plate count versus qPCR results of pathogenicity assay of germplasm bacterial isolates on *Actinidia chinensis* var. *chinensis* ‘Hort16A’ plantlets at 7 dpi. ‘Hort16A’ plantlets were flooded with 106 CFU/mL of each bacterial isolate. Bacterial growth was quantified by A) plate count at 7 days post-inoculation (CFUs/cm2), and B) ΔCt method where the Ct value of isolate-specific bacterial primers is subtracted from the Ct of plant housekeeping gene, EF1α. Black bars represent the median values, box whiskers indicate total range of four technical replicates. Compact letter display above each box indicates significant differences between bacterial isolates using Tukey’s Honestly Significant Difference (HSD) post hoc at 5%.

**Supplementary Figure 7.**
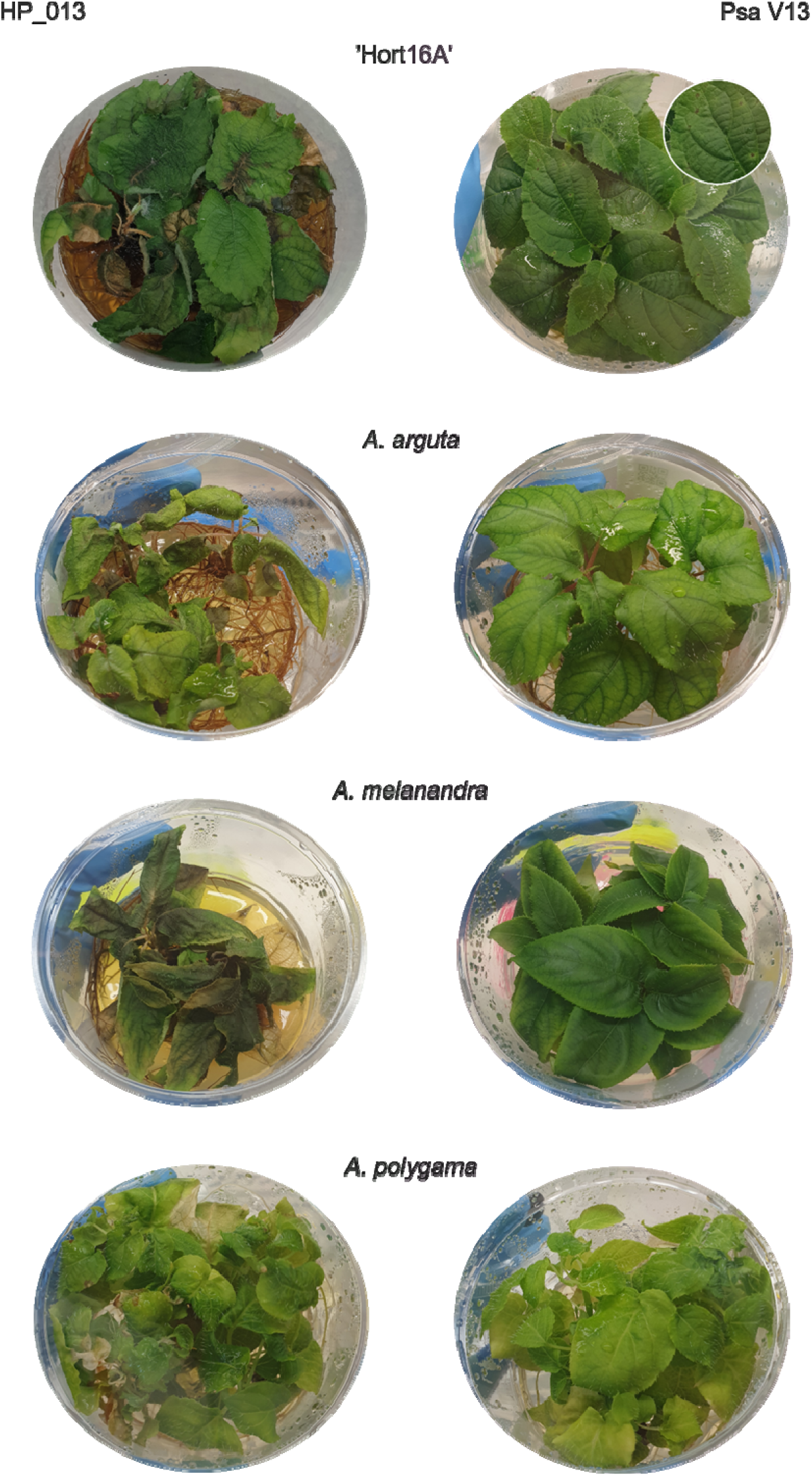
HP_013 causes tissue necrosis and agar yellowing across Actinidia hosts 7 days post-infection. Flood inoculation of tissue culture *A. arguta* AA07_03, *A. polygama* PC02_01, *A. melanandra* ME02_01, and *A. chinensis* var. chinensis ‘Hort16A’ with *Pseudomonas syringae* pv. *actinidiae* biovar 3 (Psa3) V-13 and HP_013.

**Supplementary Table 1.**
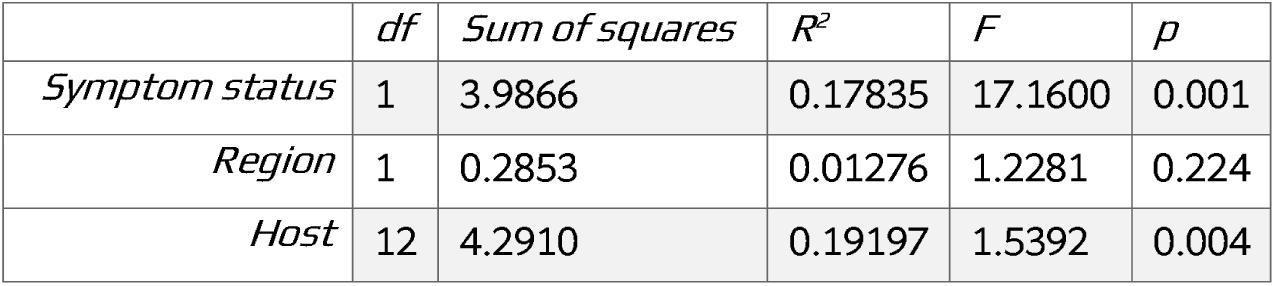
PERMANOVA results using Bray-Curtis dissimilarity matrix. Table shows the discrete variables symptom status, region, and host with their associated statistics (df = degrees of freedom, F = F-statistic, R^2^ = coefficient of determination, p = p-value). Statistics were calculated using vegan function adonis().

**Supplementary Table 2.**
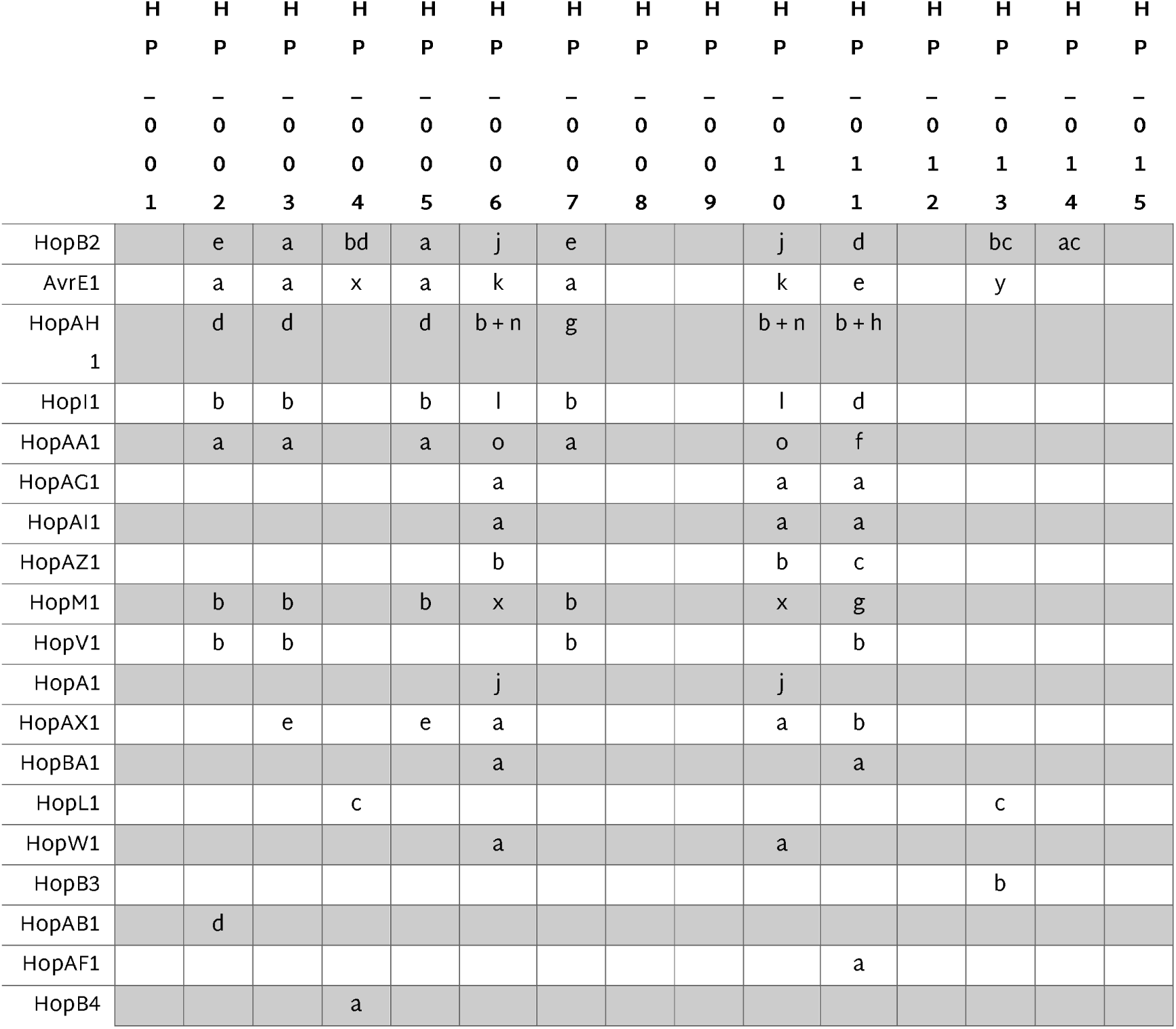
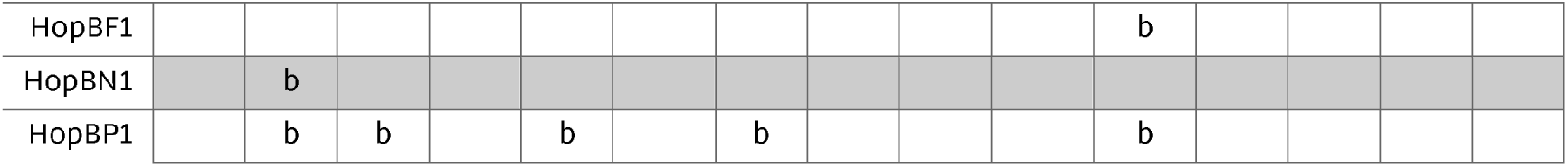
Effector families and subtypes identified in HP germplasm isolates. Isolates are noted in the header row, while the effector family is described in the left-most column. Within each cell, effector subtype is denoted with a letter.

**Supplementary Table 3.** Summary of interactions between germplasms isolates and Psa3 V-13 when co-inoculated onto *Actinidia* tissue culture plantlets. This table summarises whether the combination of the germplasm isolate or Pfm with Psa3 V-13 resulted in increased, decreased, or unaffected growth for either party. Pfm is described with the relevant host to which the result pertains.

|  |  | Psa3 V-13 |  |  |
| --- | --- | --- | --- | --- |
|  |  | Increased | Decreased | Unaffected |
| Germplasm isolate | Increased | HP_004<br>HP_013<br>HP_014<br>Pfm (in *A. arguta*) |  |  |
|  | Unaffected | HP_015 | HP_007<br>HP_009<br>Pfm (in *A. melanandra*) | HP_001<br>HP_006<br>Pfm (in *A. polygama*) |

**Supplementary Table 4.** Germplasm samples selected for amplicon sequencing of the phyllosphere bacterial community. Details are provided for each sample including sampling region, block within the germplasm, and the *Actinidia* species and variety that was sampled. Samples are given a unique code for identification, typically a number which is followed by ‘C’ for asymptomatic samples.

| Region | Block | Sample ID | Kiwifruit species |
| --- | --- | --- | --- |
| Te Puke | Block 48 | 36 | *A. hypoleuca* |
| Te Puke | Block 48 | 39 | *A. melanandra* |
| Te Puke | Block 48 | 41 | *A. melanandra* |
| Te Puke | Block 48 | 42 | *A. melanandra* |
| Te Puke | Block 48 | 44 | *A. melanandra* |
| Te Puke | Block 48 | 52 | *A. polygama* |
| Te Puke | Block 48 | 54 | *A. polygama* |
| Te Puke | Block 48 | 56 | *A. polygama* |
| Te Puke | Block 48 | 63 | *A. chrysantha* |
| Te Puke | Block 48 | 67 | *A. callosa* |
| Te Puke | Block 43 | 68 | *A. latifolia* |
| Te Puke | Block 43 | 72 | *A. rufa* |
| Te Puke | Block 43 | 75 | *A. rufa* |
| Te Puke | Block 48 | 85 | *A. glaucophylla* |
| Te Puke | Block 48 | 91 | *A. arguta* |
| Te Puke | Block 48 | 93 | *A. arguta* |
| Te Puke | Block 48 | 99 | *A. arguta* |
| Kerikeri | Block 14 | 101 | *A. arguta* |
| Kerikeri | Block 14 | 105 | *A. melanandra* |
| Kerikeri | Block 14 | 115 | *A. chrysantha* |
| Kerikeri | Block 14 | 119 | *A. latifolia* |
| Kerikeri | Block 14 | 133 | *A. guilinensis* |
| Kerikeri | Block 14 | 139 | *A. guilinensis* |
| Kerikeri | Block 3 | 155 | *A. melanandra* |
| Kerikeri | Block 3 | 162 | *A. arguta* |
| Kerikeri | Block 3 | 164 | *A. hypoleuca* |
| Kerikeri | Block 3 | 169 | *A. callosa* |
| Kerikeri | Block 3 | 177 | *A. rufa* |
| Kerikeri | Block 3 | 187 | *A. rufa* |
| Kerikeri | Block 3 | 197 | *A. arguta* |
| Kerikeri | Block 3 | 199 | *A. valvata* |
| Kerikeri | Block 3 | 207 | *A. macrosperma* |
| Kerikeri | Block 3 | 209 | *A. polygama* |
| Kerikeri | Block 3 | 211 | *A. polygama* |
| Kerikeri | Block 3 | 217 | *A. eriantha* |
| Kerikeri | Block 3 | 222 | *A. eriantha* |
| Kerikeri | Block 3 | 223 | *A. polygama* |
| Kerikeri | Block 14 | 101C | *A. arguta* |
| Kerikeri | Block 14 | 105C | *A. melanandra* |
| Kerikeri | Block 14 | 115C | *A. chrysantha* |
| Kerikeri | Block 14 | 119C | *A. latifolia* |
| Kerikeri | Block 14 | 133C | *A. guilinensis* |
| Kerikeri | Block 14 | 139C | *A. guilinensis* |
| Kerikeri | Block 3 | 155C | *A. melanandra* |
| Kerikeri | Block 3 | 162C | *A. arguta* |
| Kerikeri | Block 3 | 164C | *A. hypoleuca* |
| Kerikeri | Block 3 | 169C | *A. callosa* |
| Kerikeri | Block 3 | 177C | *A. rufa* |
| Kerikeri | Block 3 | 187C | *A. rufa* |
| Kerikeri | Block 3 | 197C | *A. arguta* |
| Kerikeri | Block 3 | 199C | *A. valvata* |
| Kerikeri | Block 3 | 207C | *A. macrosperma* |
| Kerikeri | Block 3 | 209C | *A. polygama* |
| Kerikeri | Block 3 | 211C | *A. polygama* |
| Kerikeri | Block 3 | 217C | *A. eriantha* |
| Kerikeri | Block 3 | 222C | *A. eriantha* |
| Kerikeri | Block 3 | 223C | *A. polygama* |
| Te Puke | Block 48 | 36C | *A. hypoleuca* |
| Te Puke | Block 48 | 39C | *A. melanandra* |
| Te Puke | Block 48 | 41C | *A. melanandra* |
| Te Puke | Block 48 | 42C | *A. melanandra* |
| Te Puke | Block 48 | 44C | *A. melanandra* |
| Te Puke | Block 48 | 52C | *A. polygama* |
| Te Puke | Block 48 | 54C | *A. polygama* |
| Te Puke | Block 48 | 56C | *A. polygama* |
| Te Puke | Block 48 | 63C | *A. chrysantha* |
| Te Puke | Block 48 | 67C | *A. callosa* |
| Te Puke | Block 43 | 68C | *A. latifolia* |
| Te Puke | Block 43 | 72C | *A. rufa* |
| Te Puke | Block 43 | 75C | *A. rufa* |
| Te Puke | Block 48 | 85C | *A. glaucophylla* |
| Te Puke | Block 48 | 91C | *A. arguta* |
| Te Puke | Block 48 | 93C | *A. arguta* |
| Te Puke | Block 48 | 99C | *A. arguta* |

**Supplementary Table 5.**
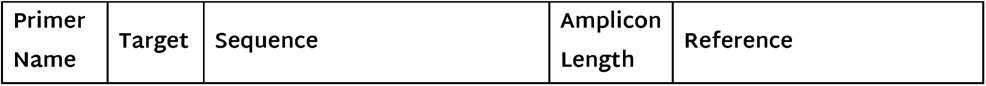
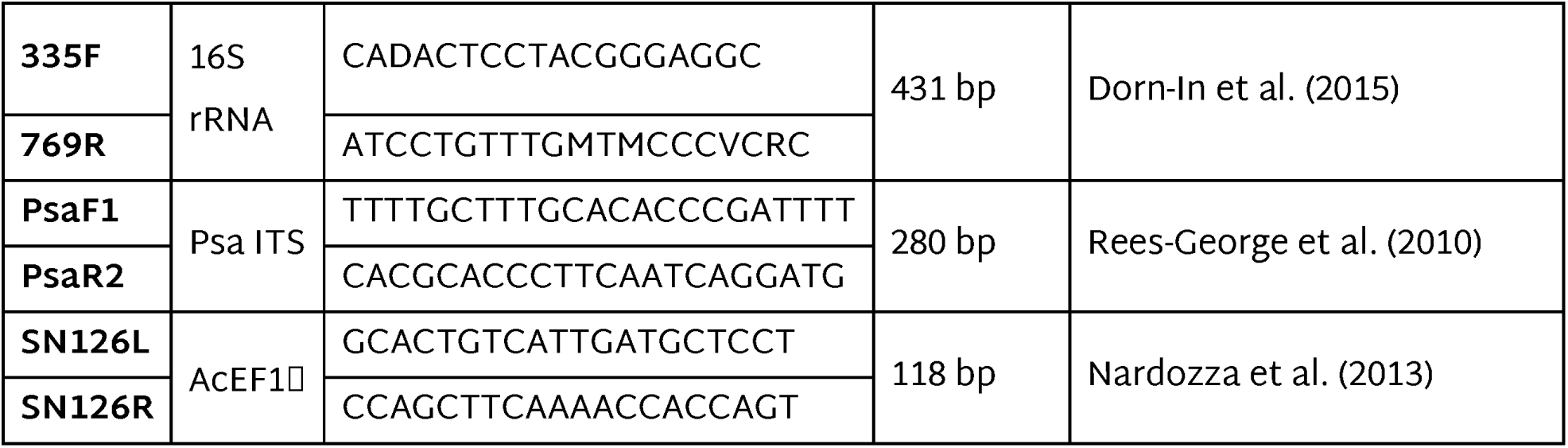
Primers used in qPCR. Primers’ common names, target, sequence, and amplicon length is listed as well as the reference from which the primers were identified.

**Supplementary Table 6.**
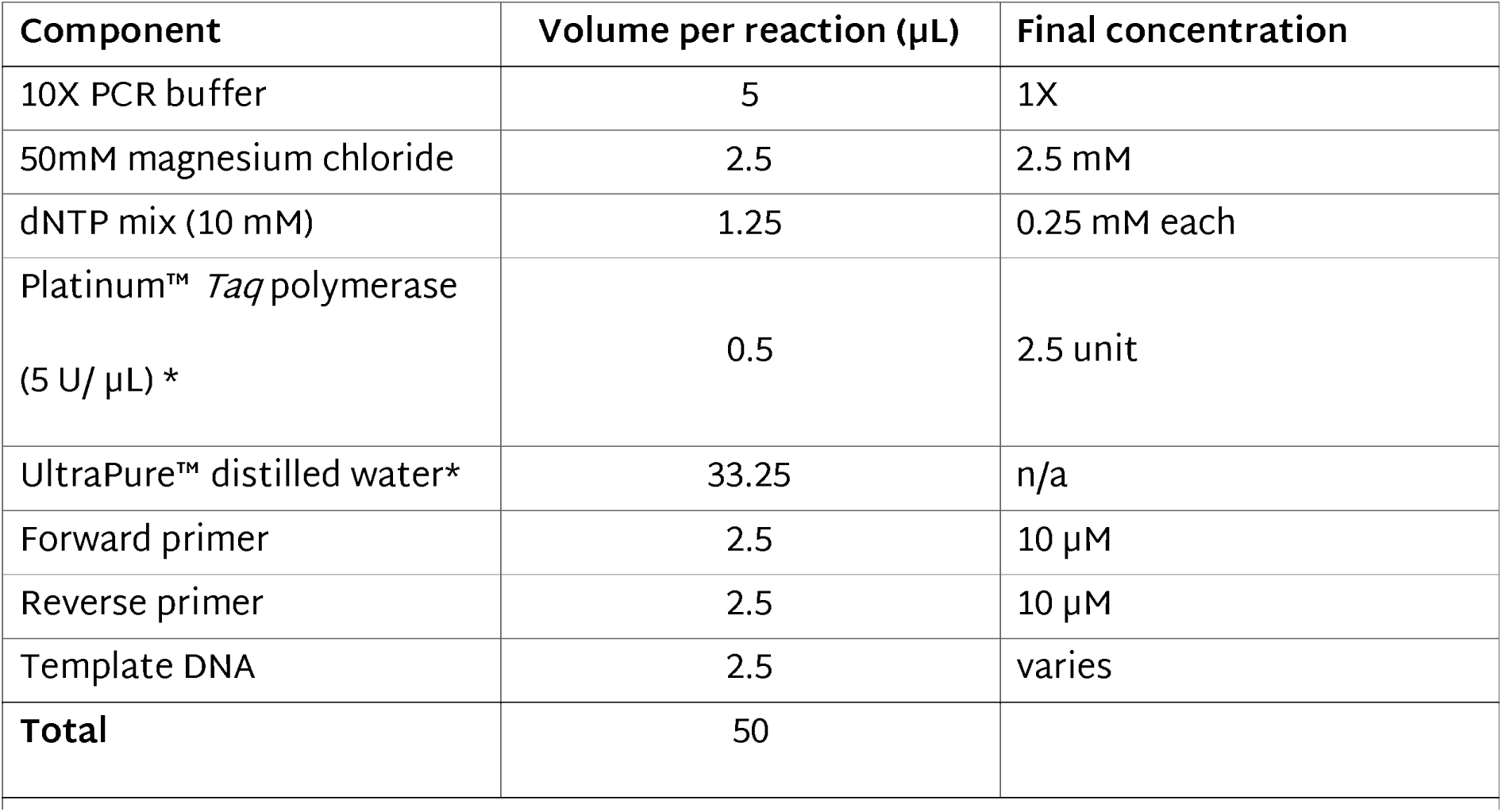
End-point PCR master-mix components. *Invitrogen (Thermo-Fisher Scientific), California, USA.

**Supplementary Table 7.** End-point PCR cycle parameters.

| Cycling step | Temperature | Time | Number of cycles |
| --- | --- | --- | --- |
| Enzyme activation | 94°C | 4 min | 1 |
| Denaturation | 95°C | 30 s | 30 or 40 (depending<br>on sample type) |
| Annealing | 57°C | 30 s |  |
| Extension | 65°C | 45 s |  |
| Final extension | 65°C | 10 min | 1 |
| Hold | 10°C | n/a | n/a |

**Supplementary Table 8.** Metadata for germplasm isolates that underwent whole genome sequencing. The original identifying number given at sampling and the isolate code and number is listed. Isolate numbers were only given to samples that were analysed for genome features. The *Actinidia* host and genotype, as well as region and block they were sampled from is also given.

| Sample ID | Isolate name | Year | Host | Region |
| --- | --- | --- | --- | --- |
| 10 | HP_001 | 2022 | *Actinidia arguta* | Te Puke |
| 12 | HP_002 | 2022 | *Actinidia arguta* | Te Puke |
| 19 | HP_003 | 2022 | *Actinidia arguta* | Te Puke |
| 33 | HP_004 | 2022 | *Actinidia arguta* | Te Puke |
| 40 | HP_005 | 2022 | *Actinidia melanandra* | Te Puke |
| 43 | HP_006 | 2022 | *Actinidia melanandra* | Te Puke |
| 44 | HP_007 | 2022 | *Actinidia melanandra* | Te Puke |
| 87 | HP_008 | 2022 | *Actinidia arguta* | Te Puke |
| 89 | HP_009 | 2022 | *Actinidia arguta* | Te Puke |
| 154 | HP_010 | 2022 | *Actinidia melanandra* | Kerikeri |
| 155 | HP_011 | 2022 | *Actinidia melanandra* | Kerikeri |
| 159 | HP_012 | 2022 | *Actinidia melanandra* | Kerikeri |
| 179 | HP_013 | 2022 | *Actinidia arguta* | Kerikeri |
| 214 | HP_014 | 2022 | *Actinidia polygama* | Kerikeri |
| 228 | HP_015 | 2022 | *Actinidia polygama* | Kerikeri |

**Supplementary Table 9.** Type III secretion system (T3SS) representative isolates. Each representative isolate was used as the reference for T3SS investigation in germplasm isolates.

| **T3SS** | **Representative isolate** | **Assembly or GenBank accession** |
| --- | --- | --- |
| **Canonical T-PAI** | *Pseudomonas syringae* pv.<br>*actinidiae*<br>ICMP 18884 | GCF_000648735.3 |
|  | *P. syringae* pv. *tomato* DC3000 | GCF_000007805.1 |
| **Alternate T-PAI** | *P. viridiflava* ICMP 3272 | GCF_001466905.1 |
|  | *P. viridiflava* PNA3.3a | AY597278.1 |
| **PG13 A-PAI** | *P. quasicaspiana* ICMP 23753 | GCF_025643095.1 |
|  | *P. syringae* UB246 | GCA_000452865.1 |
| **PG2c A-PAI** | *P. syringae* pv. *syringae* 642 | GCA_000177515.1 |
| **R-PAI** | *P. syringae* pv. *phaseolicola* 1448a | GCF_000012205.1 |
| **S-PAI** | *P. viridiflava* RMX3.1b | AY597283.1 |
|  | *P. viridiflava* LCMA8 | GCF_000834695.1 |

**Supplementary Table 10.**
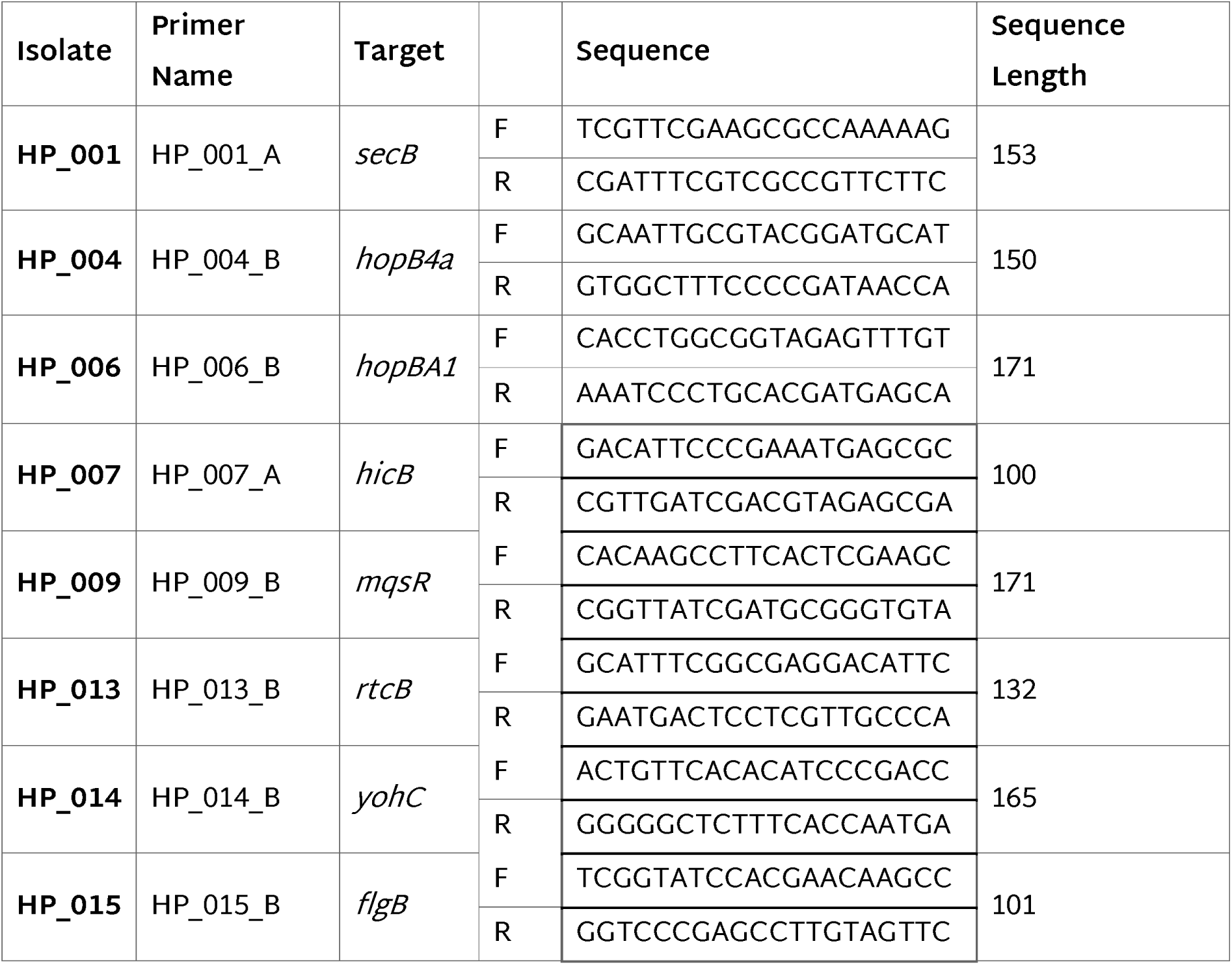
Isolate-specific primers used for quantifying bacterial growth in *planta* using qPCR. Targeting primers were developed for each germplasm isolate, which is outlined in the following table including the isolate, the primer name, target gene, forward and reverse sequences, and the expected sequence length.

